# RNA/DNA Binding Protein TDP43 Regulates DNA Mismatch Repair Genes with Implications for Genome Stability

**DOI:** 10.1101/2024.05.16.594552

**Authors:** Vincent E. Provasek, Albino Bacolla, Suganya Rangaswamy, Manohar Kodavati, Joy Mitra, Issa O. Yusuf, Vikas H. Malojirao, Velmarini Vasquez, Gavin W. Britz, Guo-Min Li, Zuoshang Xu, Sankar Mitra, Ralph M. Garruto, John A. Tainer, Muralidhar L. Hegde

## Abstract

TDP43 is an RNA/DNA binding protein increasingly recognized for its role in neurodegenerative conditions, including amyotrophic lateral sclerosis and frontotemporal dementia (FTD). As characterized by its aberrant nuclear export and cytoplasmic aggregation, TDP43 proteinopathy is a hallmark feature in over 95% of ALS/FTD cases, leading to the formation of detrimental cytosolic aggregates and a reduction in nuclear functionality within neurons. Building on our prior work linking TDP43 proteinopathy to the accumulation of DNA double-strand breaks (DSBs) in neurons, the present investigation uncovers a novel regulatory relationship between TDP43 and DNA mismatch repair (MMR) gene expressions. Here, we show that TDP43 depletion or overexpression directly affects the expression of key MMR genes. Alterations include MLH1, MSH2, MSH3, MSH6, and PMS2 levels across various primary cell lines, independent of their proliferative status. Our results specifically establish that TDP43 selectively influences the expression of MLH1 and MSH6 by influencing their alternative transcript splicing patterns and stability. We furthermore find aberrant MMR gene expression is linked to TDP43 proteinopathy in two distinct ALS mouse models and post-mortem brain and spinal cord tissues of ALS patients. Notably, MMR depletion resulted in the partial rescue of TDP43 proteinopathy-induced DNA damage and signaling. Moreover, bioinformatics analysis of the TCGA cancer database reveals significant associations between TDP43 expression, MMR gene expression, and mutational burden across multiple cancers. Collectively, our findings implicate TDP43 as a critical regulator of the MMR pathway and unveil its broad impact on the etiology of both neurodegenerative and neoplastic pathologies.

## Introduction

Transactive response DNA binding protein of 43 kDa (TDP43) is a nuclear protein that regulates the expression of a wide range of genes, including its transcripts (1–3). This highly conserved RNA/DNA-binding protein is furthermore linked to multiple pathological processes, including neurodegeneration and cancer (4–6). TDP43 is expressed in a variety of tissues, with particularly high levels in the brain and spinal cord, and is essential for the normal development and function of the nervous system (7). RNA processing is a complex and tightly regulated process that involves numerous RNA-binding proteins to facilitate appropriate RNA processing. TDP43 plays a central role in several aspects of RNA processing, including alternative splicing, mRNA stability, and transport (8–10). For example, TDP43 binds to the 3’-untranslated region (UTR) of target mRNAs to regulate their stability and localization (11, 12). Additionally, TDP43 interacts with various splicing factors to coordinate efficient intron removal from target transcripts (13–16). The precise mechanisms by which TDP43 regulates RNA processing are not completely understood, yet several studies have provided important insights into this process. One speculation could be that TDP43 may act as a scaffold protein that brings together different components of the RNA processing machinery, thereby facilitating their interactions and coordination (11). In addition, TDP43 facilitates the recruitment of other regulatory proteins to target mRNA, influencing either their stabilization or degradation (8, 17, 18). The specific mechanisms that functionally link TDP43 and DNA repair have also been shown in the context of neurodegenerative disease. We have reported that TDP43 is a critical scaffolding factor required for the DNA double-strand break (DSB) repair (19). However, given the extensive role of TDP43 in RNA metabolism, we questioned whether it may contribute to DNA repair in a more global aspect via its effects on the processing of DNA repair gene transcripts. Here, we discover a previously unknown role of TDP43 in modulating the expression of MMR genes.

MMR is a critical DNA repair pathway responsible for at least a 10-fold increase in the replication fidelity of replicating cells (20). Deficient MMR (dMMR) is classically associated with neoplastic pathologies, most notably Lynch Syndrome (21–23). The MMR family consists of six core members - MLH1, MSH2, MSH3, MSH6, PMS2, and PMS1. These components function as heterodimers that recognize mispaired or otherwise aberrant nucleic acid substrates and initiate their excision and replacement (24, 25). The recognition heterodimers, MutSα (MSH2-MSH6) and MutSβ (MSH2-MSH3), interact with the MutLα (MLH1-PMS2) heterodimer to begin the repair process; this is followed by several other DNA repair factors, including RPA, PCNA, exonuclease 1 and DNA polymerase δ, to excise and re-synthesize damaged sequences (26–30). Importantly, defects in any of these factors, particularly MutSα or MutLα, are sufficient to disrupt the entire MMR machinery and thus lead to increased mutational load and tumorigenesis.

While the consequences of dMMR on cancer development are well known, other functions of MMR are less understood. One of these functions involves the role of MMR at the intersection between DNA repair and the DNA damage response (DDR) signaling pathway. The relationship between MMR and DDR has been reviewed elsewhere (31–35). However, the importance of this interaction is evinced by the requirement of MMR for chemotherapy-induced (CTX) cancer cell killing, particularly following select platinum-based, and DNA alkylator agents. Reports of CTX resistance in colon, endometrium, and CNS cancers, among others, demonstrate a correlation with dMMR and subsequently reduced cell killing (36–40). The function of MMR-induced DDR activation is at the center of this apparent paradox. Specifically, it was shown that DNA damage-induced DDR activation did not occur without an active MMR pathway, and cells did not undergo subsequent apoptosis (41). Overabundance of MMR has likewise been attributed to negative effects on genome integrity. Limited reports show that overexpression of certain MMR factors (e.g., MLH1 or PMS2) induces a hypermutable phenotype *in vivo* (42, 43). Moreover, others have shown that overexpression of MSH2, irrespective of its repair capability, is sufficient to induce cell killing, possibly through direct activation of apoptosis via specific domains within the protein (44). Perhaps it should come as no surprise that the MMR repair system could indeed act in a pro-instability manner, given its documented role in facilitating somatic hypermutation during immunoglobulin class switch recombination (33, 35, 45–47). In any case, the ability of the MMR system to either repair or induce mutations depends on cellular context and expression regulation.

The question of MMR activity and regulation takes on renewed importance in the neurodegeneration field, where less is known regarding the role of MMR in the nondividing cell. Early reports have suggested that MMR activity in the context of non-dividing cells conferred a mutagenic effect, possibly due to a loss of strand discrimination signals (48, 49). In neurons, the function of MMR is even less clear. At least one study suggests that neuronal MMR would be limited to the repair of deamination products, including 5-methyl cytosine to thymine, or cytosine to uracil (50). More recently, however, there has been renewed interest in the role of MMR in neurons, especially within the context of neurodegeneration. Recent reports have indicated that MMR may contribute to the expansion and downstream clinical severity of trinucleotide repeat (TNR) mutations like those observed in Huntington’s disease (51–56). Taken together, there remain many significant questions regarding MMR function, particularly in non-dividing cells. One clear theme, however, is that the expression of MMR proteins appears to exist in a delicate balance that, if disturbed, may have important consequences for cellular function, genomic integrity, and degenerative disease.

In this study, we shed light on at least one mechanism maintaining this balance by reporting for the first time that TDP43 regulates the expression of MMR genes. We showed that TDP43 exerts this role across multiple cell lines and animal models. We furthermore traced the mechanism of this effect by showing that TDP43 acts to regulate the pre-mRNA splicing and transcript stability of at least two key MMR transcripts, such as *MLH1* and *MSH6*. By using animal models of TDP43-associated ALS (ALS-TDP43) and CNS tissues of Guamanian ALS patients, we discovered that the expression of MMR genes is indeed altered at the protein level. Moreover, we showed that similar changes observed using in vitro models of TDP43 pathology also increased the levels of DNA damage markers and impaired cellular response to oxidative DNA damage. Finally, we showed that TDP43-mediated expression of MMR genes also extends to various cancers. By using extensive bioinformatic analysis, we found informative correlations between TDP43 and MMR gene expression, with concomitant associations with patient survival. Collectively, these findings link TDP43 to the active regulation of the MMR pathway through the modulation of key MMR protein levels. Consequently, TDP43 dysregulation may contribute to the pathogenesis of two major age-related disease conditions of our time - neurodegeneration and cancer.

## Methods

### Cell Culture

Human neuroblastoma SH-SY5Y cells (ATCC) or HEK293 (ATCC) cells were cultured in DMEM/F12 media (Gibco) or DMEM High-glucose (Gibco) with 10% Fetal Bovine Serum (Sigma) and 1% Penicillin-Streptomycin (Sigma) at 37 °C with 5% CO_2_. SH-SY5Y cells were differentiated with Retinoic acid (RA) (10 μM) and BDNF (50 ng/mL) in DMEM/F12 and 1% FBS for 4 to 6 days. WT human neural stem cells were (Jackson Labs) maintained on Geltrex LDEV-Free in Stem Pro medium (Gibco). Terminally differentiated neurons were derived from NPCs (Jackson Lab) WT (JIPSC1000), TDP43-Q331K (JIPSC1066), TDP43-A315T and TDP43-M337V, according to established methods (106). Briefly, NPSCs were plated at 2.5–5 ×10^4^ cells/cm^2^ in 60 cm dish coated with Poly-D-Lysine and Laminin and initially maintained using StemPro media (Thermo Fisher) before transitioning to Neural Differentiation Media consisting of Neurobasal Medium (Gibco), 2% B-27 Serum-Free Supplement (Gibco), 2 mM GlutaMAX-I Supplement (Gibco). On day 7, dibutyryl cAMP (Sigma Cat #0627) was added at 10 µM concentration for 3 days to complete neural differentiation. For human iPSC to motor neuron differentiation, control iPSC clones KYOU-DXR0109B (201B7) (ATCC) were initially plated on a 60-cm dish, then passaged to a T-25 flask with neuronal basic medium (mixture of 50% Neurobasal medium and 50% DMEM/F12 medium, with N2 and B27 supplements without vitamin A), following collagenase type IV digestion. After 2 days incubating with 5 μM ROCK Inhibitor (Y-27632, RI, Merck Millipore), 40 μM TGF-β inhibitor (SB 431524, SB, Tocris Bioscience), 0.2 μM bone morphogenetic protein inhibitor (LDN-193189, LDN, Stemgent), and 3 μM GSK-3 inhibitor (CHIR99021, CHIR, Tocris Bioscience), suspended cell spheres were then incubated with a neuronal basic medium containing 0.1 μM retinoic acid (RA, from Sigma) and 500 nM Smoothened Agonist (SAG, from Merck Millipore) for 4 days. Cells were then incubated for 2 days in a neuronal basic medium containing RA, SAG, 10 ng/ml Brain-derived neurotrophic factor (BDNF, Peprotech), and 10 ng/ml Glial cell-derived neurotrophic factor (GDNF, Peprotech). Cell spheres were then dissociated with a neuronal basic medium containing trypsin (0.025%)/DNase in the water bath for 20 min at 37 °C. Then they were pipetted into single cells with the medium containing trypsin inhibitor (1.2 mg/mL). After cell counting, a defined number of cells were seeded into 20 μg/mL Laminin (Life Technologies) -coated dishes or chamber slides and incubated for 5 days in a neuronal basic medium containing RA, SAG, BDNF, GDNF, and 10 μM DAPT, then incubated for 2 days in a neuronal basic medium containing BDNF, GDNF, and 20 μM Inhibitor of γ-secretase (DAPT, Tocris Bioscience). For MN maturation, cells were kept for over 7 days in a medium containing BDNF, GDNF, and 10 ng/ml ciliary neurotrophic factor (CNTF, Peprotech). The same method was followed for culturing ALS patient-derived iPSC line (a kind gift from Dr. Ludo Van Den Bosch) and differentiating to NPSCs and iMNs (107).

The ALS patient-derived iPSC lines carrying TDP43 mutations at G287S, G298S, A382T with C9orf72 (a kind gift from the collaborators Drs. Ludo Van Den Bosch and Philip Van Damme at the Stem cell institute, KU Leuven) were used for the successive differentiation into the respective NPSC and iMN lines, following the method described elsewhere (107).

### Transfection, Plasmids, and siRNAs

SH-SY5Y and HEK293/T cells were transfected with siRNA using Lipofectamine 3000 (Thermo Fisher) per the manufacturer’s instructions. NSCs were transfected using Lipofectamine Stem Reagent (Thermo Fisher) per the manufacturer’s instructions. Gene KD was achieved using Silencer Select siRNAs (Thermo Fisher) for Control (Cat #4390843), TDP43 (Cat #4390771), MLH1 (AM51331), and MSH2 (AM16708). FLAG-tagged TDP43 WT or ΔNLS plasmids in the pCW backbone (Addgene #50661) were generated by cloning the target coding DNA sequences (CDS) by PCR amplification and restriction digestion of the vector and insert at NheI and BamHI sites (41).

### Inducible Cell Lines

Doxycycline (Dox)-inducible TDP43 WT and ΔNLS models were generated in the SH-SY5Y background, as described elsewhere (5, 19). During the induction period, cells were cultured in a reduced FBS (1%) containing DMEM/F12 media, facilitating the protein mislocalization and aggregation processes in the cytosol.

### Protein Isolation and Immunoblotting (IB)

Total cell lysates from SH-SY5Y, HEK293, NPSC lines, mouse cortical tissues, and human CNS tissues were extracted with RIPA lysis buffer (Millipore) containing the protease/phosphatase inhibitor cocktail (Thermo Fisher). Mouse and human tissue were first flash-frozen in liquid nitrogen and homogenized by mortar and pestle. After mixing with RIPA lysis buffer, total homogenates were probe sonicated at low amplitude, 3 times for 5 sec each, on ice. After centrifugation at 16000xg for 15 min at 4°C, supernatants containing the soluble protein fraction were decanted and quantified using the Bradford assay. The remaining pellet containing the insoluble protein fraction was resuspended in 2% SDS in PBS buffer and probe sonicated twice before gel electrophoresis. Protein isolates were mixed with protein loading buffer containing 100 mM DTT reducing agent and denatured for 5 min at 95°C before loading onto 4–12% Bis-Tris NuPAGE precast gels (Thermo Fisher) for electrophoresis. Following transfer to nitrocellulose membrane and incubating with primary and secondary antibodies, the protein signal was detected using the LI-COR XF imaging system and analyzed using Empiria Imaging Software (LI-COR). Where applicable, total protein stains of transferred blots were performed using Revert Total Protein Stain (LI-COR) and used to normalize target protein expression. Primary antibodies used: MLH1 (Cell Signaling #4256), MSH2 (Cell Signaling #2017), MSH3 (Protein Tech #22393-1-AP), MSH6 (Protein Tech #18120-1-AP), PMS2 (ABClonal #A6947), GAPDH (Cell Signaling #2118), γH2AX (Cell Signaling #2577), H2AII (Cell Signaling #2578), TDP43 (Protein Tech #10782-2-AP), FLAG (Sigma #F1804), cleaved-caspase-3 (Cell Signaling #9664), and PARP-1 (Cell Signaling #9542).

### Cell Death/Survival Assay

Treated or untreated HEK cells exposed to 6-thioguanine (6-TG; 10 μM) or methylmethanesulfonate (1 mM) were seeded in 96-well ELISA plates (Corning) in triplicate. Cultures were incubated 48 h post-treatment. MTT assay was performed following the manufacturer’s protocol (TREVIGEN). Briefly, 10 μL of 5 mg/mL MTT reagent (Sigma) was added to each well and incubated for 24 h, then 100 μL of 10% SDS in PBS detergent (Thermo Scientific) was added. Reagents on plates were gently mixed by rotational agitation for 10 min before measuring the absorbance at 570 nm using a microplate reader (Bio-Rad, 680 XR).

### Mouse Models

The endogenous Tdp43ΔNLS mouse model was generated by CRISPR/Cas9-mediated knock-in of NLS-deleted Exon3 of the mouse *Tardbp* gene flanked by loxP sequences within the intronic region of the gene. Conditional expression of Cre recombinase driven by the Ubc promoter (motor neuron-specific) in the bigenic Cre::Tdp43ΔNLS mice induced the expression of murine Tdp43ΔNLS, resulting in the manifestation of human ALS-TDP43-type proteinopathy in the central nervous system (https://doi.org/10.21203/rs.3.rs-3879966/v1). Brain tissue samples from a moderately overexpressing murine Tdp43 ALS mouse model were kindly provided by the laboratory of co-author Dr. Zuoshang Xu (70). Mice’s ages ranged from 94 to 396 days. Animals were anesthetized using isoflurane administered through a SomnoSuite Low Flow Anesthesia System (Kent Scientific), then transcardially perfused with ice-cold phosphate-buffered saline (PBS, Thermo Scientific). Brain tissues were dissected from the cranium and snap-frozen in liquid nitrogen. Brain tissues from five subjects of ages ranging from 94 d to 296 d for each of the following groups were used for analysis for a total of 20 mice, including the control-male, control-female, transgenic-male, and transgenic-female groups.

### Protein Isolation from Human Decedent Tissues

Autopsied cortex and spinal cord tissue samples of Guamanian ALS and age-matched controls from the same ethnic background were acquired from the Binghamton University Biospecimen Archive. The clinical characteristics of the decedents are described in Supplemental Table S6. Snap-frozen cortical tissue specimens were first ground into powder using liquid nitrogen in a mortar and pestle. For total protein extraction, about 20 mg of tissue powder from each sample was lysed using 1x RIPA buffer, added with cocktail protease and phosphatase inhibitors (Thermo Scientific). Samples were incubated on ice for 30 min and subsequently treated with 10 sec-pulse sonication three times at 5 amplitudes on ice. Samples were then centrifuged at 13000 rpm at 4°C for 15 min three times to remove most of the brain fats. Finally, the soluble and insoluble fractions of each sample were collected in separate tubes. The insoluble pellet was then resuspended in 2% SDS buffer (2% SDS, 50 mM Tris-HCl, pH 7.4). Protein concentration was determined using the Bradford assay (Sigma), according to the manufacturer’s instructions.

### Comet Assay

Alkaline comet assay was performed using SH-SY5Y cells treated with siControl or siTDP43 for 96 h as well as TDP43ΔNLS or WT cells induced with doxycycline for 72 h, according to the manufacturer’s protocol (TREVIGEN). Briefly, cells were mixed with low-melt agarose (TREVIGEN) and placed on comet assay slides (TREVIGEN), followed by overnight incubation in lysis buffer at 4°C in the dark and single-cell electrophoresis at 21 V for 30 min in alkaline electrophoresis buffer (TREVIGEN) under manufacturer-recommended conditions. Slides were finally subjected to staining with SYBR® Gold for visualization of genomic DNA under an AXIO Observer inverted microscope (Carl Zeiss).

### Immunofluorescence (IF)

IPSC-derived iMNs were cultured in 8-well chamber slides (Millicell EZ slides, Millipore) and fixed with 4% paraformaldehyde (PFA) in phosphate-buffered saline (PBS) for 30 min, then permeabilized with 0.2% Tween-20 in PBS for 15 min at room temperature. Samples were blocked with 3% BSA in PBS for 1 h. Primary antibody incubation was done with anti-TDP43 (Protein Tech #10782-2-AP) and anti-PMS2 (ABClonal #A6947) antibodies overnight at 4°C in 1% BSA solution, followed by washing three times before staining with fluorophore-tagged corresponding secondary antibodies for 1 h at room temperature in 1% BSA solution. Slides were then washed three times and nuclei were counterstained with DAPI. Images were captured with the AXIO Observer fluorescence microscope (Carl Zeiss).

### RT2-Profiler Assay

Complementary DNA (cDNA) was synthesized using total RNA isolated from siTDP43 or siControl-treated HEK293 cells. Samples were analyzed by SYBR green-based quantitative real-time (qRT) RT2 PCR arrays for evaluating expressions of a pre-defined panel of DNA damage response (DDR) genes (Qiagen #PAHS-042Z). The 96-well RT2 profiler plate contained primers for 84 DDR genes, 5 housekeeping genes, and 3 negative controls (Supplemental **Fig. S1A**).

### qRT-PCR and End-Point PCR

Approximately 2×10^6^ cells were collected and centrifuged at 4°C at 2000 rpm for 3 min for extraction of RNA. Total RNA was isolated using Trizol (Thermo Scientific), according to manufacturer’s instructions. RNA purity and quantification were determined using a NanoDrop 2000 spectrophotometer (Thermo Scientific). Total RNA was reverse transcribed into cDNA using the SuperScript Vilo Kit (Thermo Scientific). The qRT-PCR amplification was performed in triplicate using the ABI 7500 (Applied Biosystems) using the PowerUp SYBER Green Master Mix (Thermo Scientific). *HPRT* and *GAPDH* housekeeping genes were utilized as internal controls. Primers were purchased from Sigma and are shown in Supplemental **Tables S2** and **S3**. The relative expression of each validated gene was determined using the 2^-ΔΔCt^ method. Two-sided student’s t-test was performed and results with p<0.05 (*) were statistically significant.

### Transcript Splicing Analysis

Complementary DNA (cDNA) was synthesized from total RNA isolated from control and TDP43 KD HEK293 cells and used for MLH1 and MSH6 splicing analysis using the End-Point PCR assay. To determine whether intronic sequences were included in the transcripts of TDP43 KD cells, primers were designed to anneal to two consecutive exon sequences surrounding one intronic sequence. The intervening cDNA sequence was then amplified using LongAmp Taq polymerase. Primers for each gene region are shown in Supplemental **Table S3**. Amplicons were then subject to electrophoresis on a 2% agarose gel (Sigma) and stained with ethidium bromide (Fisher Scientific) before visualization using the LI-COR XF Imaging System.

### Minigene assay

A minigene assay was conducted to further validate the role of TDP43 in regulating the splicing of MMR genes using the RHCglo reporter plasmid in HEK293 cells with or without TDP43 KD (64). Genomic regions encompassing exon 17, intron 17, and exon 18 were amplified from HEK293 cells and cloned at BamHI and XhoI sites of the RHCglo plasmid. The resulting positive clones were transfected into both control and TDP43 KD cells. At 48 h post-transfection, the cells were harvested and total RNA was extracted. cDNA was synthesized from 1 μg of RNA, and PCR was performed using primers from the RHCglo plasmid that flank the cloning site (Supplemental Table S4). The PCR products were analyzed on a 2% agarose gel.

### RNA Stability Assay

The stability of target RNA transcripts was assessed as previously described (108, 109). SH-SY5Y cells underwent two treatments with Silencer Select siRNA against TDP43 (Thermo #4392420) on days 6 and 7 after seeding cells on the plate. On day 8, control cells were harvested, while the experimental cells were pre-treated with 10 μM of α-amanitin (Sigma) before harvesting at 8 h, 12 h, and 24 h timepoints. cDNA was prepared from total RNA and quantified by qRT-PCR. Normalization of C_t_ values was performed using 18s rRNA as the internal control. Fold-change at each timepoint was calculated, relative to the control, and values were fitted to a one-phase decay plot to determine the half-life of each transcript under consideration using GraphPad Prism software.

### Bioinformatic Analysis

TCGA RNA-seq transcriptomic data and clinical patient data were obtained using the R package TCGA-assembler v. 2 (110) and processed with custom scripts to obtain gene expression data in both tumors and matched controls, as well as Kaplan-Meier survival curves (111). The tumor abbreviations used were: ACC, adrenocortical carcinoma; BLCA, bladder urothelial carcinoma; BRCA, breast invasive carcinoma; CESC, cervical squamous cell carcinoma and endocervical adenocarcinoma; CHOL, cholangiocarcinoma; COAD, colon adenocarcinoma; DLBC, lymphoid neoplasm diffuse large B-cell lymphoma; ESCA, esophageal carcinoma; GBM, glioblastoma multiforme; HNSC, head and neck squamous cell carcinoma; KICH, kidney chromophobe; KIRC, kidney renal clear cell carcinoma; KIRP, kidney renal papillary cell carcinoma; LAML, acute myeloid leukemia; LGG, brain lower grade glioma; LIHC, liver hepatocellular carcinoma; LUAD, lung adenocarcinoma; LUSC, lung squamous cell carcinoma; MESO, mesothelioma; OV, ovarian serous cystadenocarcinoma; PAAD, pancreatic adenocarcinoma; PCPG, pheochromocytoma and paraganglioma; PRAD, prostate adenocarcinoma; READ, rectum adenocarcinoma; SARC, sarcoma; SKCM, skin cutaneous melanoma; STAD, stomach adenocarcinoma; TGCT, testicular germ cell tumors, THCA, thyroid carcinoma; THYM, thymoma, UCEC, uterine corpus endometrial carcinoma; UCS, uterine carcinosarcoma; UVM, uveal melanoma.

Data were plotted using the R libraries “ggplot2”, “ggpubr”, “extrafont”, “dplyr”, “survival”, and “survminer” and SigmaPlot (https://systatsoftware.com/sigmaplot/), and further edited in Canvas (https://www.canvasgfx.com/products/canvas-x-draw). Functional annotations of gene sets were performed with DAVID (https://david.ncifcrf.gov/); ARCHS4 (https://maayanlab.cloud/archs4/) was used to retrieve “Predicted biological processes (GO)”, “Predicted pathways (KEGG)”, and “Most similar genes based on co-expression” from large sets of RNA-seq data for *TARDB* and key MMR-related genes. For correlations between gene expression and mutational burden in TCGA, for each gene and each tumor type, patients were divided into two groups, a g_high group with RNA-seq data above population mean, and a g_low group with RNA-seq data below population mean. TCGA patients codes for the g_high and g_low groups were then intersected with the same patients codes from the Catalogue Of Somatic Mutations In Cancer (COSMIC, https://www.sanger.ac.uk/tool/cosmic/; CosmicMutantExport.tsv file, version 92) to retrieve the curated list of simple somatic mutations, single bases substitutions and small insertions and deletions, in the tumor samples, and the differences in mutational loads (log10 of number of mutations) between g_high and g_low were assessed with t tests. Matching COSMIC patient codes were available for 24 of the 33 TCGA tumor types. Heatmap of statistical differences in mutational loads was generated with Heatmapper (http://heatmapper.ca). Values used for the heatmap were the -log10 p-values for h_high>g_low, and log10 p-values for g_high<g_low. Scores were reported as z-score, i.e. the number of standard deviations each value distanced from the mean across the 24 tumor types. C++ codes for t-tests and linear regressions were used in pipeline custom scripts.

For the TP43 CLIP-seq analysis, File GSM998871_MP41.BED.gz containing the mouse CLIP-seq Tdp43 results was downloaded from https://www.ncbi.nlm.nih.gov/geo/query/acc.cgi?acc=GSM998871. Genes and coordinates of Tdp43-bound RNA were extracted using Bash commands. DNA sequences of the corresponding Tdp43-bound RNA were extracted using TwoBitToFa (https://hgdownload.soe.ucsc.edu/admin/exe/linux.x86_64/) on the mm9.2bit mouse genome downloaded from ftp://hgdownload.cse.ucsc.edu/goldenPath/mm9/bigZips/. TGn or CAn (depending on strand occupied by Tdp43) Tdp43 binding motifs within the RNA CLIP-seq sequences were retrieved using the Bash “grep” command, choosing a minimum length of 6 units, i.e. grep -E “(TG){6,}|(CA){6,}”.

## Results

### TDP43 Regulates the Expression of DNA Mismatch Repair Genes

An exploratory investigation of TDP43’s effects on mRNA expression of DNA repair genes was first conducted using an RT2-Profiler assay (Qiagen) with HEK293 cells, treated with either control (siControl) or TDP43-directed (siTDP43) small interfering RNAs. By downregulating the TDP43 expression to approximately 50% of control levels (**Fig. 1A**), we found significant changes in the expression of multiple DNA repair protein families (**Fig. 1B-C**, Supplemental **Table S1**). Of those families identified, MMR factors MLH1, MSH3, MSH6, and PMS2 exhibited a greater than two-fold decrease in their gene expressions. These results were confirmed by qRT-PCR (**Fig. 1D** and Supplemental **Table S2)** and immunoblotting (IB) analyses (**Fig. 1E)** using HEK293 cultures with TDP43 knockdown (KD). We next questioned how the MMR genes’ expressions might change upon TDP43 overexpression (OE). Notably, moderate OE of wildtype (WT) TDP43 in differentiated neuron-like SH-SY5Y cells was sufficient to boost the expression of key MMR factors markedly beyond their basal levels (**Fig. 1F**).

**Figure 1.**
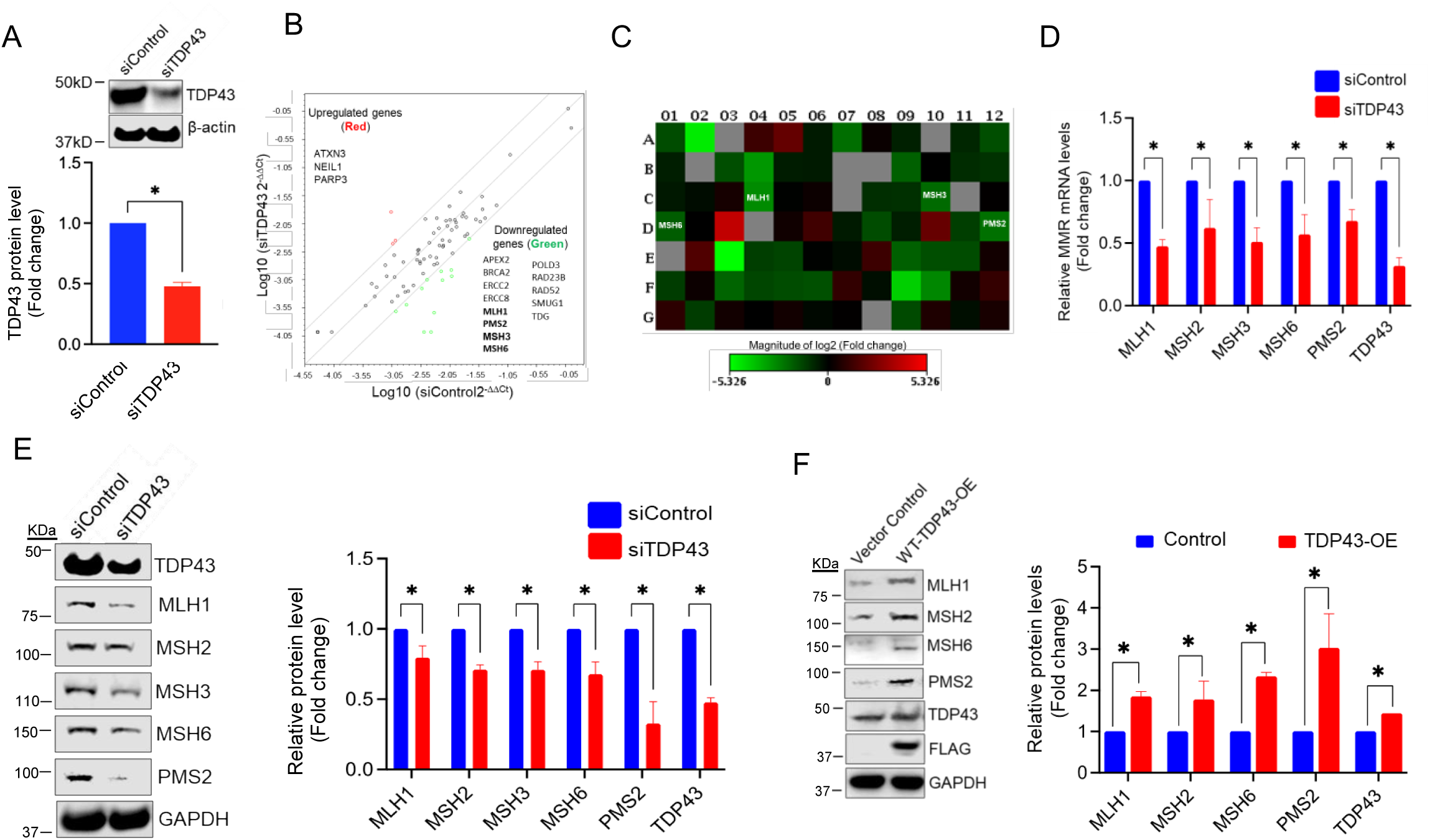
TDP43 Modulates the Expression of DNA Mismatch Repair (MMR) factors. (**A**) Western blot analysis of TDP43 knockdown (KD) in siControl or siTDP43 treated HEK293 cells. β-actin served as the loading control. Quantitation of relative TDP43 protein levels is shown in the bottom. (**B**) Scatter plot of RT2-Profiler DNA Repair Expression Array representing genes in siControl or siTDP43-treated cells that display >2-fold difference in mRNA expressions compared with that of control cells. Red, green, and black circles indicate upregulated genes, downregulated genes, and nonregulated genes, respectively. (**C**) Heatmap highlighting altered expressions of MMR and other DNA repair genes in siControl versus siTDP43 cells. Red, green, and black squares indicate upregulated genes, downregulated genes, and unaffected genes, respectively. (**D**) Histogram showing the relative mRNA transcript levels of MMR factors in siControl or siTDP43-treated HEK293 cells. (**E**) Western blot (WB) images illustrate the comparison of MMR protein expressions between the siControl and siTDP43-treated HEK293 cells. And the quantitation of relative protein levels in fold changes. (**F**) WB analysis of the expression of indicated MMR factors with or without TDP43 OE in SH-SY5Y cells. Quantitation of relative protein levels in fold changes, normalized to the GAPDH level shown in histogram (right). Significance values (p-values) are as follows: p >0.5 (ns), p <0.5 (*). Error bars indicate mean ± SEM, from three independent biological replicates.

Given these results, we questioned whether TDP43 KD could decrease the expression of MMR genes in induced pluripotent stem cell (iPSC)-derived neural progenitor stem cells (NPSCs). Using IB (**Fig. 2A-B**) and immunofluorescence (IF) (**Fig. 2C**) approaches, we observed that siTDP43-mediated TDP43 KD caused decreased expression of MLH1, MSH3, MSH6, and PMS2 in NPSCs. This further clarified that TDP43 is necessary and sufficient for regulating the expression of MMR factors in iPSC-derived NPSCs.

**Figure 2.**
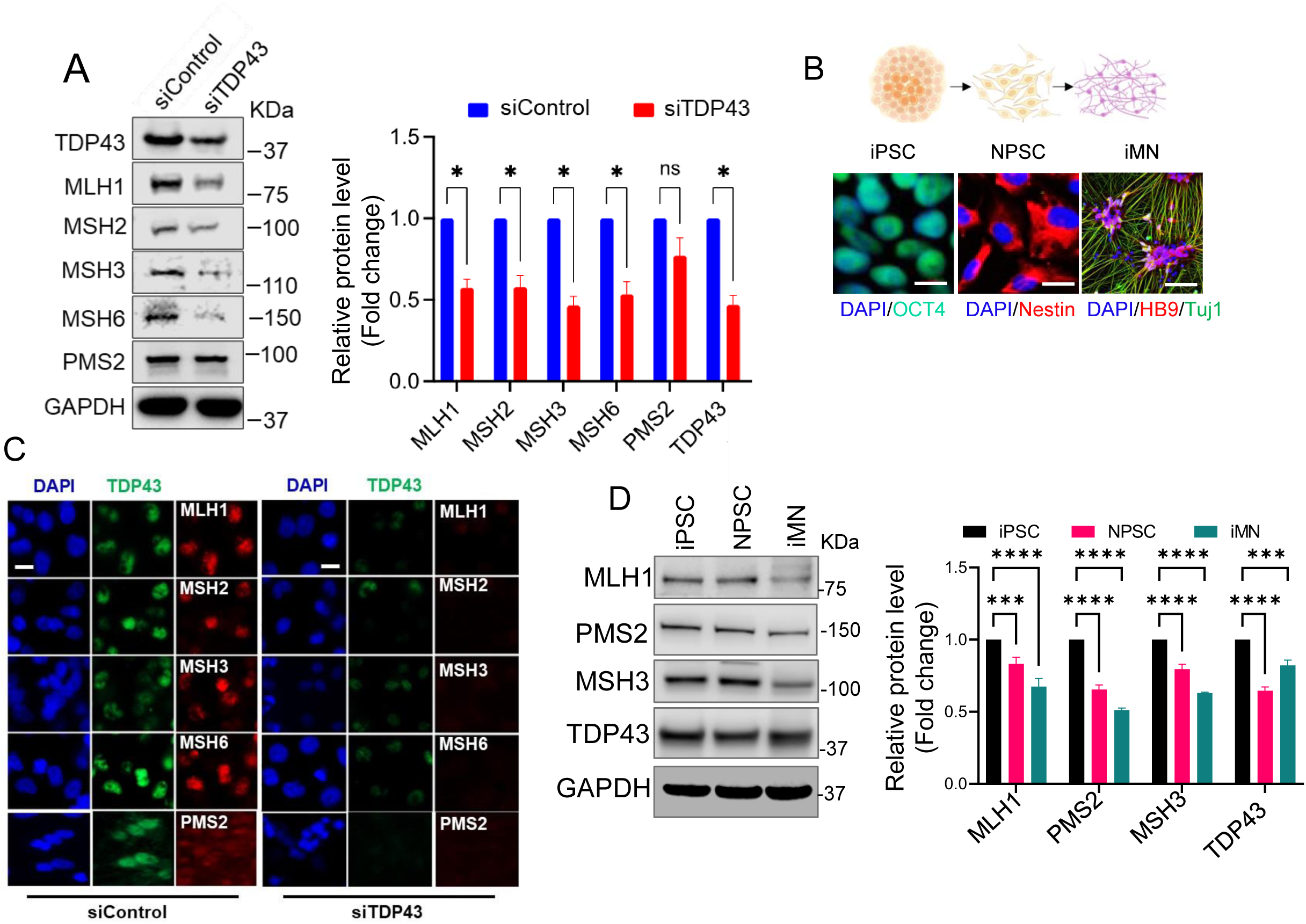
Differential expressions of MMR genes in iPSC-derived neuronal cells and their regulation by TDP43. (**A**) WB images and histogram illustrating a quantitative comparison of MMR expression in control and siTDP43-treated human pluripotent stem cell (iPSC)-derived neural lineage progenitor stem cells (NPSCs). Quantitation of protein levels normalized to that of GAPDH. (**B**) Schematic of iPSC, NPSC, and terminally differentiated motor neurons (iMN) utilized in this study. (**C**) Immunofluorescence (IF) images revealing the expression of TDP43 (green); and MLH1 (red), MSH2 (red), MSH3 (red), MSH6 (red) and PMS2 (red), and nuclear DNA (DAPI-blue) after siTDP43-mediated KD of TDP43 in iPSC-derived iMNs. Significance values (p-values) are as follows p >0.5 (ns), p <0.5 (*), p <0.01 (**), p <0.001 (***) **D**) WB images show the expression of select MMR factors across the following states of cellular differentiation: iPSC, NPSC, iMN.. Error bars indicate mean ± SEM from three independent experiments.

We next assessed the expression levels of MMR proteins in both dividing and non-dividing neuronal cells, such as NPSCs and iPSC-derived induced motor neurons (iMNs), respectively. To do so, iPSCs were differentiated to iMNs via the NPSC stage, as illustrated in **Fig. 2D**. IB analysis of MMR factors in NPSCs and iMNs revealed that levels of MLH1, MSH2, and MSH3 were significantly reduced in the iMN stage compared to those in the NPSC stage. These results underscore the significance of TDP43-mediated regulation of MMR protein expression in DNA repair processes regardless of the cell cycle status.

### TDP43 Depletion and Mutation Alter Transcriptional Processing of MMR Transcripts

TDP43 interacts with several splicing factors to ensure the efficient processing of pre-mRNAs into mature mRNAs as well as prevent pathological alternative splicing events of its target transcripts (14, 15, 57, 58), as illustrated in **Fig. 3A**. Disruption of this process can lead to the retention of intronic sequences, dysregulating the nuclear export, cytosolic degradation, or synthesis of alternative protein products. Moreover, perturbations of this process have been reported to contribute significantly to the neuropathology of multiple diseases, especially ALS and FTD (4, 12, 59–61).

**Figure 3.**
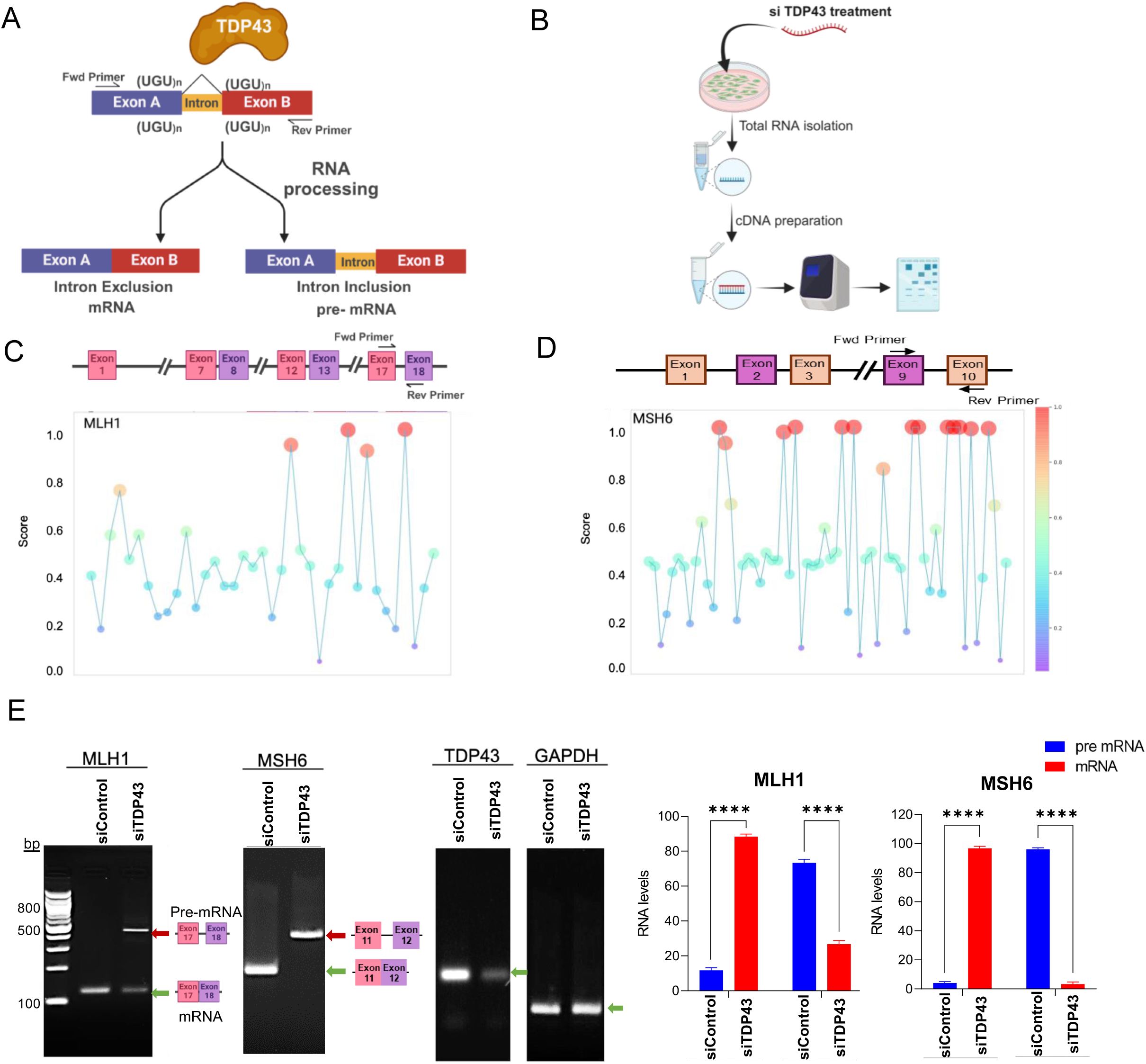
TDP43 depletion alters transcriptional processing of specific MMR factors. (**A-B**) A conceptual representation illustrating TDP43’s anticipated role in pre-mRNA processing and a schematic of experimental design/ workflow employed to characterize TDP43-induced splicing of MMR transcripts by quantitating relative pre-mRNA versus mRNA transcript levels. (**C-D**) *In silico* computational analysis (RBPSuite) MLH1 and MSH6 sequences identified multiple putative TDP43 binding sites across each transcript. These identified sequences guided the primer design, to amplify exon-intron-exon junctions for post-cDNA synthesis amplification. (**E**) PCR amplification of selected regions of the MLH1 amplifying the region between Exon 17—Exon 18, and MSH6 amplifying the region between Exon9—Exon10 transcripts derived from cDNA prepared from HEK293 cultures treated with either siControl or siTDP43, histograms showing fold change of band intensity representing pre mRNA and fully spliced mRNA. Significance values (p-values) are as follows: p >0.5 (ns), p <0.0001 (****). Error bars indicate mean ± SEM from three independent experiments.

Given these functions, we asked whether TDP43 might regulate MMR expression by affecting the processing of MMR transcripts. To this end, we first probed the splicing patterns of two key MMR gene transcripts, namely *MLH1* and *MSH6*, by qRT-PCR analysis of total RNA from HEK293 cells with or without TDP43 KD (**Fig. 3B-E** and Supplemental **Table S3**). To inform our selection of transcript regions, we employed two complementary *in silico* approaches: RBPSuite (62) and RBPMap (v1.2) (63). Primers (Supplemental **Table S2**) were designed to PCR amplify selected Exon-Intron-Exon regions with relatively high predictive scores for TDP43 binding of each target gene (**Fig. 3C-D**), such that the size of the PCR product would reflect the presence or absence of intronic DNA sequences. Based on these predictions, we selected regions between Exon 17 and Exon 18 of *MLH1* and Exon 8 and Exon 9 of *MSH6* as potential TDP43 splicing targets. We then performed siRNA-mediated TDP43 KD and measured the resulting mRNA levels by qRT-PCR. In the case of MHL1 and MSH6, TDP43 KD caused a greater than 50% decrease in the fully spliced mRNA product and a concomitant increase of at least one predicted unspliced variant (**Fig. 3E**). Evaluations of other MMR gene (MSH3 and PMS2) transcripts displayed either no changes in splicing patterns or sporadic decreases of fully spliced transcripts, respectively (Supplemental **Fig. S1A-B**). To further validate the role of TDP43 in regulating splicing of MMR genes, we performed a minigene splicing assay using the RHCglo reporter system in TDP43 KD HEK293 cells, following the protocol described elsewhere (64). A genomic fragment spanning exon 17, intron 17, and exon 18 of the *MLH1* gene was PCR amplified from HEK293 cells and cloned into the RHCglo plasmid at BamHI-XhoI sites. We observed efficient splicing of intronic sequences in siControl cells, as evident by ∼700 bp band, while intron retention was dramatically high in siTDP43-treated cells (Supplemental **Fig. S1C-D,** and **Table S4**). Following these analyses, we further validated the sequence-specific binding of TDP43 to MMR gene transcripts by analyzing a previously reported mouse Tdp-43 CLIP-seq dataset, retrieved from the GEO dataset (GSM998871). Supplemental **Table S5** and **Fig. S1E** show the number of TDP43 binding motifs [(UG)n] (65, 66) in the MMRG transcripts and their genomic locations. As demonstrated earlier that the expression of MMR genes is reduced in matured and/or differentiated neurons, although the frequency of TDP43 binding sites is slightly higher in the MMRG group than in the low-expression gene (LEG) group (N=20), there was no statistically significant difference in values (Supplemental **Fig. S1E**). In contrast, highly expressed genes (HEG; N=20) exhibited the presence of ∼6 times more TDP43 binding motifs per kb of genomic region of MMRG. In summary, perturbation of the TDP43 expression level can cause multi-modal dysregulations to the mRNA processing of MMRGs, and altered binding of TDP43 to its binding sites is likely to induce pathological alternatively spliced variants of its target genes, such as *MLH1* and *MSH6*.

### TDP43 Regulates the Stability of MLH1 and MSH6 Transcripts

TDP43 is shown to modulate the stability of the processed transcripts (67, 68). To identify this effect on MLH1 and MSH6 transcripts, we measured the influence of TDP43 on the stability of these transcripts. To accomplish this, we utilized a modified version of a previously described transcript stability protocol (69). Briefly, we used RT-PCR to measure the relative abundance of each transcript in total RNA samples isolated from nondividing neuronal cell cultures treated with scrambled control or TDP43 siRNAs at set timepoints after exposure to α-amanitin, a potent RNA polymerase II inhibitor (Schematically shown in **Fig. 4A**). We then fit the fold change data over time to a one-phase decay curve, which was then used to compute the approximate half-life of the transcripts. Surprisingly, we observed that the rate of MLH1 and MSH6 transcript degradation increased compared to controls during the first twelve hours of treatment (**Fig. 4B-C**). The loss of TDP43 resulted in a 38% decrease in the half-life of MLH1 transcripts and a 56% decrease in the half-life of MSH6 transcripts (**Fig. 4E**). Plots of mRNA transcript relative fold expression that were used to calculate the transcript half-life are also shown in Supplemental **Fig. S2**. Given the autoregulatory function of TDP43 on its expression, we also measured the abundance of TDP43 transcripts over time (3). Expectedly, we observed a 23% increase in the half-life of TDP43 transcripts in its KD cell cultures (**Fig. 4D**). In summary, these data indicate that TDP43 promotes the stability and persistence of MLH1 and MSH6 transcripts. This suggests that any condition that disrupts TDP43 homeostasis may also have direct effects on the MMR gene regulation.

**Figure 4.**
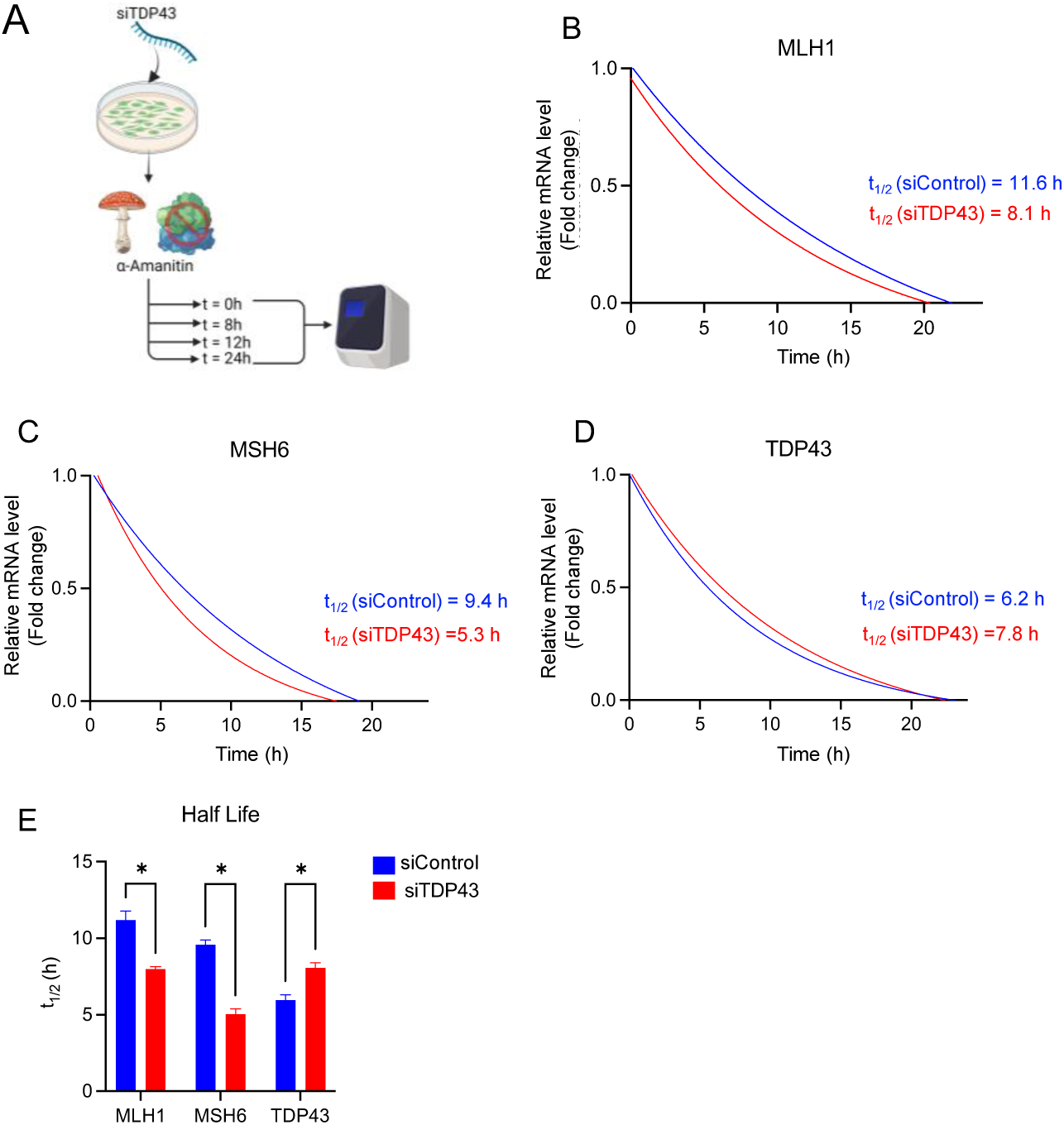
TDP43 Regulates the Stability of MLH1 and MSH6 Transcripts *in vitro*. (**A**) A schematic showing the experimental procedure for measuring the transcript half-life using nondividing neuronal cultures and the RNA polymerase II inhibitor α-amanitin. (**B-D**) Representative one-phase decay plots derived from mRNA transcript fold change expression between control and TDP43 siRNA treated cell cultures (**E**) Computed half-life (t_1/2_) values are provided for each experimental group. Significance values (p-values) are as follows: p >0.5 (ns), p <0.5 (*). Error bars indicate mean ± SEM from three independent experiments.

### TDP43 Pathology Disrupts the MMR Protein Levels *in vivo*

To determine whether MMR expression was altered within the context of neurodegenerative diseases, we utilized two distinct murine models of ALS-TDP43 (70). The first murine model utilized a CRISPR-FLeX system to allow tetracycline-inducible expression of a Tdp43 mutant lacking its nuclear localization sequence (Tdp43ΔNLS) under the ubiquitin-C (UBC) promoter (**Fig. 5A**) (71). This model recapitulates TDP43 mislocalization without increasing the total TDP43 level, thus isolating the effects of mislocalization itself on downstream pathways. In this model, total protein lysates of cortical brain samples showed significant increases in expression of Msh2 and Msh3, with trends toward increased expression of Msh6 (**Fig. 5B-C**). Unexpectedly, we did not observe clear increases in Mlh1 level despite increased levels of Tdp43. We attribute this increase in Tdp43 expression to a disruption in the autoregulatory function of Tdp43, where mislocalization of Tdp43 within the cytosol results in a relative loss of nuclear Tdp43 with subsequent loss of autoregulation, which was consistent with previous findings (1, 3). We suspect that the resulting increase in Tdp43 expression is endogenous in origin and may work to bind and facilitate MMR expression. To further test this hypothesis and confirm whether the increased murine Tdp43 expression was sufficient to increase MMR expression, we utilized a second transgenic model of ALS-TDP43 with moderate overexpression of murine Tdp43 under the major prion protein (PrP) promoter (**Fig. 5D**). This model involves CNS-specific moderate TDP43 OE, which leads to secondary mislocalization and neurodegeneration. In this model, we observed minor sex-specific differences in MMR expression. Both Mlh1 and Msh3 expressions were upregulated in the brain cortices of males, whereas females showed upregulation of Mlh1, Msh2, and Msh3 (**Fig. 5E-G** and Supplemental **Fig. S3**).

**Figure 5.**
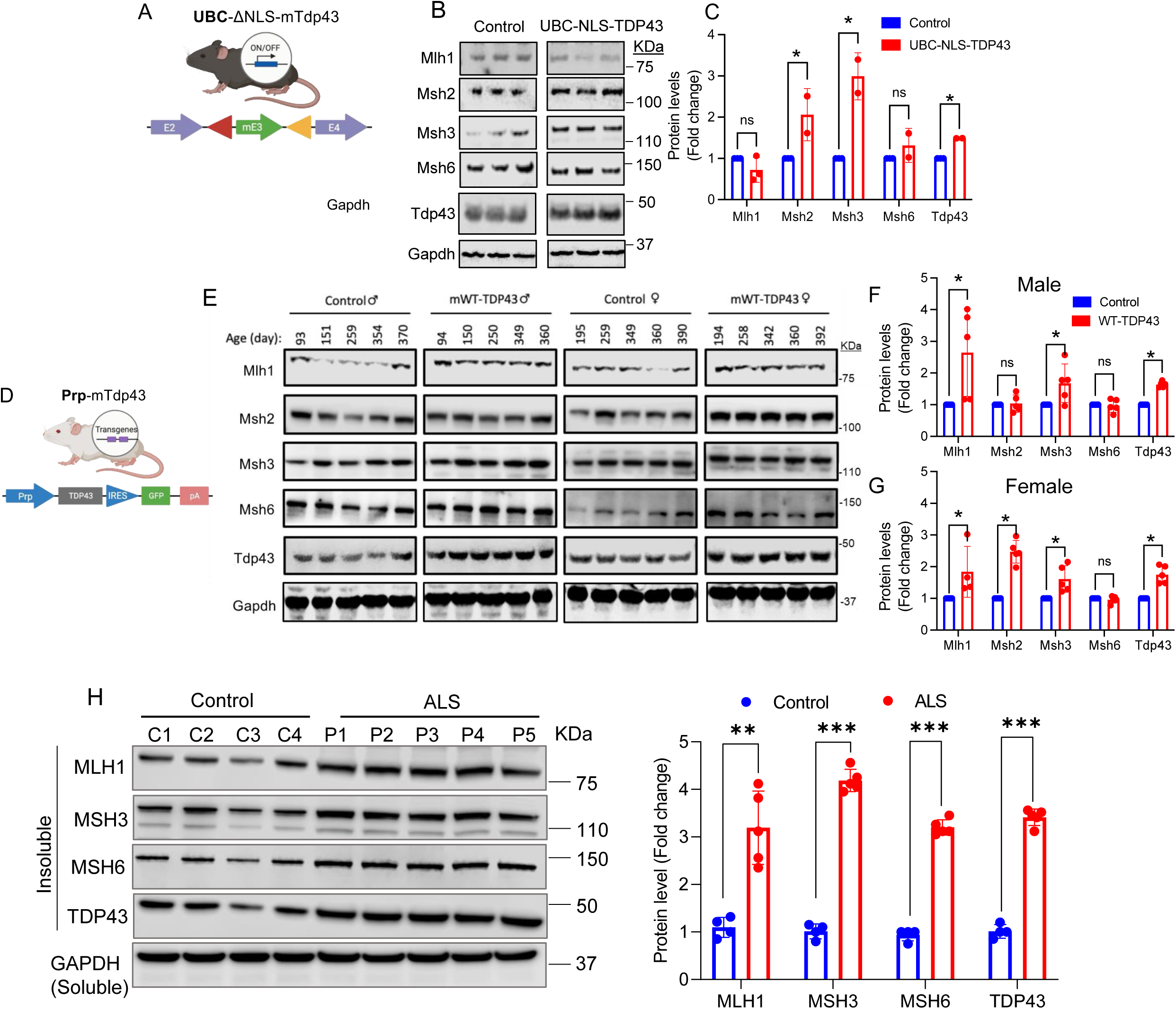
Mouse models of TDP43-ALS display an altered MMR expression phenotype. **A**) Schematic of the construct used to generate TDP43-ΔNLS expression under the UBC promoter in C57BL6 mice. **B**) WB of cortical brain tissue lysate from three control and three transgenic (Tg) mice are shown. **C**) Histograms showing fold change in immunoblot band intensity of averaged Tg samples (red) relative to control (blue) indicate significant increases in MSH2, MSH3, and trends toward increased MSH6 in TDP43-ΔNLS mice. **D**) Schematic of the construct used to generate moderate overexpression of WT murine TDP43 (mTDP43) under the PrP promoter in FVB/NJ mice. **E**) Representative immunoblots of cortical brain tissue lysate from control and Tg mice are shown for male and female mice. **F-G**) Histograms of fold change in immunoblot band intensity of averaged Tg samples relative to control indicate significant increases in MLH1, MSH2, and MSH3 expression. **H**) WB and quantitative histogram of insoluble protein isolates from control and Guam ALS-affected human post-mortem CNS tissue. Significance values (p-values) are as follows: p >0.5 (ns), p <0.5 (*), p <0.01 (**), p <0.001 (***). Error bars indicate standard error mean ± SD, derived from at least three biological and technical replicates.

Given these results in mouse models of ALS-Tdp43, we next examined whether MMR protein expression was altered in the post-mortem CNS tissues of Guamanian ALS-Parkinsonian (Guam ALS) affected patients. IB analysis of insoluble protein fractions of total protein from Guam ALS brain samples revealed consistent increases in the detection of MLH1, MSH3, and MSH6 proteins relative to non-neurological controls (**Fig. 5H** and Supplemental **Table S6**). In summary, these data demonstrate that TDP43 modulates the level of key MMR factors *in vivo*.

### Depletion of MMR Factors Rescues TDP43 Pathology-Associated DNA Damage

The roles of MMR toward DDR signaling and apoptosis have been previously described (32, 34, 41, 72). To determine if loss of TDP43 confers similar functional effects on DNA damage-induced cell killing as direct MMR KD, we employed the MTT assay. HEK293 cell cultures were treated with 10 μM 6-thioguanine (6-TG) for 24 h followed by a 24 h recovery period before adding MTT reagent to culture media. MMR-deficient cultures were achieved via simultaneous siRNA KD of both MLH1 and MSH2 (siMMR). In line with previous reports, MMR deficiency prevented 6-TG mediated cell killing while combined MLH1 and MSH2 overexpression (OE-MMR) had the opposite effect. Importantly, TDP43 KD also mitigated 6-TG cell killing while TDP43 overexpression enhanced cell killing compared to controls (**Fig. 6*A***). These results indicate that TDP43-mediated changes in MMR expression may also influence cellular response to DNA damage.

**Figure 6.**
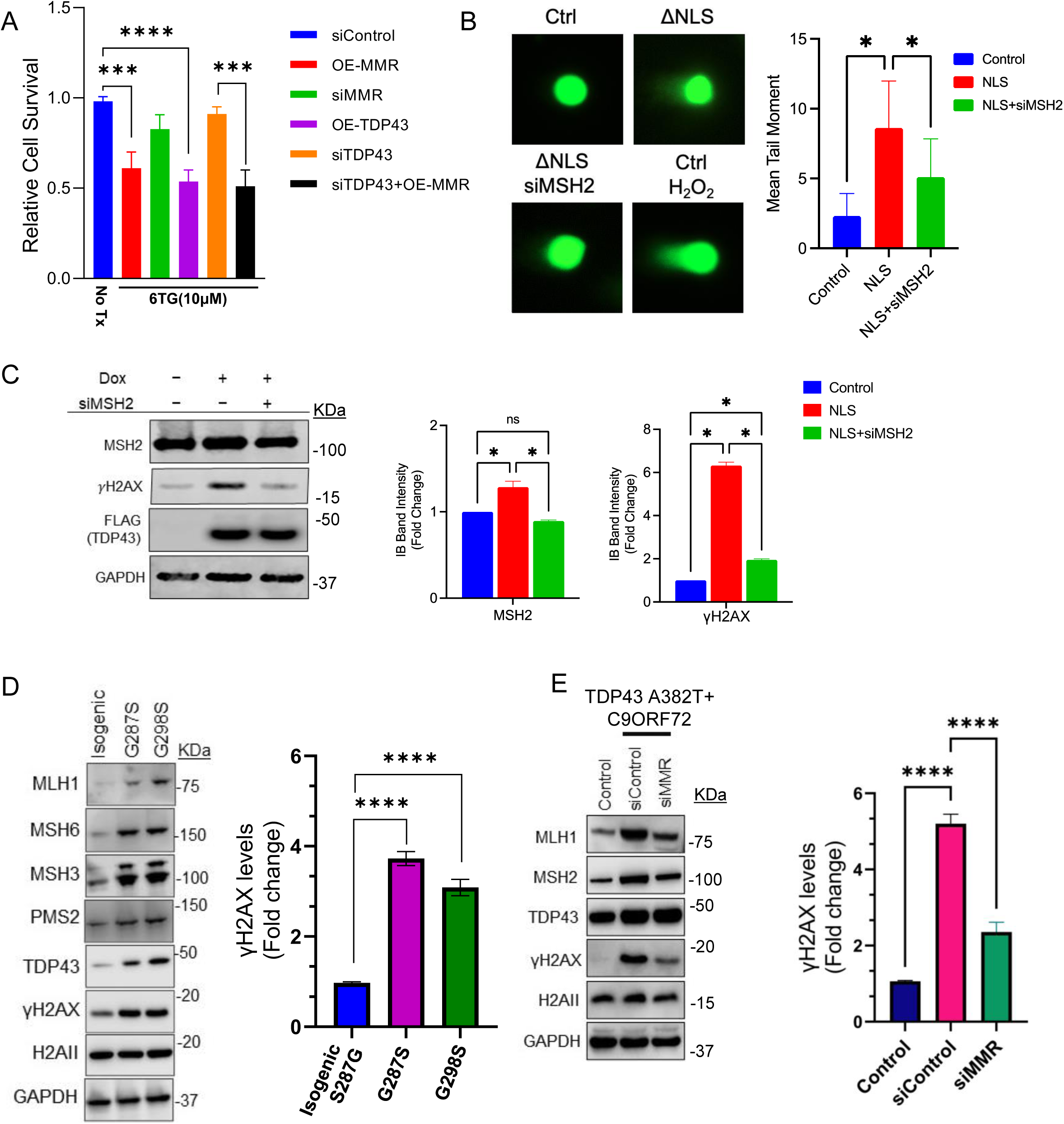
MMR expression modulates TDP43 proteinopathy-induced DNA damage. **A)** MTT assay showing cellular viability of HEK293 cell cultures treated with 10 mM (also change in figure) 6-thioguanine (6-TG) relative to non-treated controls. Experimental groups included scrambled siRNA control (siControl), siRNA knockdown TDP43 (siTDP43), siRNA knockdown of MLH1 and MSH2 (siMMR), overexpression (OE) of wildtype (WT) TDP43 (OE-TDP43), and MMR factors MLH1 and MSH2 (OE-MMR), and siTDP43 with OE-MMR. **B)** Alkaline single-cell gel electrophoresis (Comet Assay) in TDP43ΔNLS (NLS) showing increased levels of DNA damage relative to controls, which is partially rescued by siRNA knockdown of MSH2. The histogram shows quantification of the mean olive tail moment across each condition. **C)** IB analysis with quantitation histograms of NLS expressing cells (Dox 2.5 mg/mL for five days) shows increased levels of DNA damage marker *γ*H2AX that decrease following siRNA knockdown of MSH2. **D)** IB analysis of MMR and DNA damage markers with indicated antibodies in two distinct ALS patient-derived iPSC-derived NPSC lines carrying TDP43 mutations at G287S and G298S, and an isogenic mutation-corrected control line (S287G). GAPDH served as the loading control for total protein and H2AII as the control for γH2AX level normalization. Quantitation of γH2AX levels. Error bars indicate mean ± SD, derived from at least three biological and technical replicates. **E)** IB analysis of control or TDP43 A382T+C9ORF72 patient cells with or without siMMR (MLH1+MSH2). GAPDH was the loading control. And quantitation of γH2AX levels. Significance values (p-values) are as follows: p >0.5 (ns), p <0.5 (*). p <0.001(****). Error bars indicate standard error mean ± SD, derived from at least three biological and technical replicates.

We next questioned whether MMR may contribute to TDP43 proteinopathy-induced DNA damage. To this end, we utilized a cell model of TDP43-ALS. Cultures of SH-SH5Y cells carrying a Dox-inducible plasmid expressing TDP43ΔNLS were used to recapitulate TDP43 proteinopathy. After differentiation, cultures were induced (Dox 2.5 mg/mL) for five days to yield significant cellular pathology, increased basal levels of DNA damage, and DDR activation. To identify potential MMR-mediated DNA damage, we utilized alkaline single-cell gel electrophoresis (Comet assay) as a sensitive measure of SSBs and DSBs. Interestingly, we observed significant increases in DNA damage in TDP43ΔNLS cell lines, which was partially mitigated by KD of MSH2 (**Fig. 6B**). IB analysis was also performed using samples of these cultures to assess γH2AX levels as a marker of DNA damage (**Fig. 6C**). The reason we chose MSH2 as the primary target for downregulation is because MSH2 forms essential complexes with either MSH3 (MutSβ) or MSH6 (MutSα), depending on the nature of DNA mismatch, and sensitizes the MMR machinery toward damage response and repair (35, 73–75). Therefore, MSH2 KD is likely to lead to the collapse of the overall MMR pathway.

Results from this experiment showed that TDP43ΔNLS expression alone caused significant increases in DNA damage marker γH2AX and evidence of pathologic TDP43 aggregates, as identified by bands at 43 kDa and 35 kDa (Supplemental **Fig. S4A**). Expectedly, MSH2 expression was also upregulated. Importantly, MSH2 KD appeared to diminish TDP43 proteinopathy-induced DNA damage, as indicated by lower γH2AX expression (**Fig. 6C**).

Given the effect of MSH2 KD on basal levels of TDP43 proteinopathy-induced DNA damage, we hypothesized that the MMR system would also affect DNA repair capacity. To this end, we tested the ability of MMR proteins MLH1 and MSH2 to regulate DNA repair activities in terms of γH2AX levels in an ALS patient-derived NPSC line transfected with or without siControl or siMMR (siMLH1 + siMSH2), revealing a three-fold decrease in γH2AX levels in siMMR-treated cells compared to siControl cells (**Fig. 6D-E**). Cells treated with 100 nM glucose oxidase (GO) for 45 min were allowed to recover for 6 h after treatment. IB analysis of these cell extracts showed not only an increase in markers of DNA damage and DDR signaling immediately after the GO challenge but also that such increases persisted in TDP43ΔNLS-expressing cells compared to controls after the recovery period. Again, KD of MSH2 appeared to improve cellular recovery from GO-induced damage, as shown by decreased levels of phosphorylated H2AX (γH2AX) and ATM (Supplemental **Fig. S4B**). These results align with previous reports describing the role of MMR in activating the DDR and underscore the regulatory role of TDP43 proteinopathy in this process.

### TDP43 Expression Correlates with the Expression of MMR Genes in Cancer

Finally, we questioned whether TDP43-mediated regulation also extended to the cancer disease context. To learn if TDP43 expression was linked to that of MMR genes in vivo, we examined transcriptomic profiles of tumors by analyzing RNA-seq data from TCGA database. First, we compared the mRNA abundance of *TARDBP* between tumor types and matched controls for which at least 10 normal samples were available (15 tumor-control pairs total), which showed that *TARDB* was overexpressed in 60% (9 total) of cases, not changed in 3 cases and downregulated in 3 cases (**Fig. 7A**).

**Figure 7.**
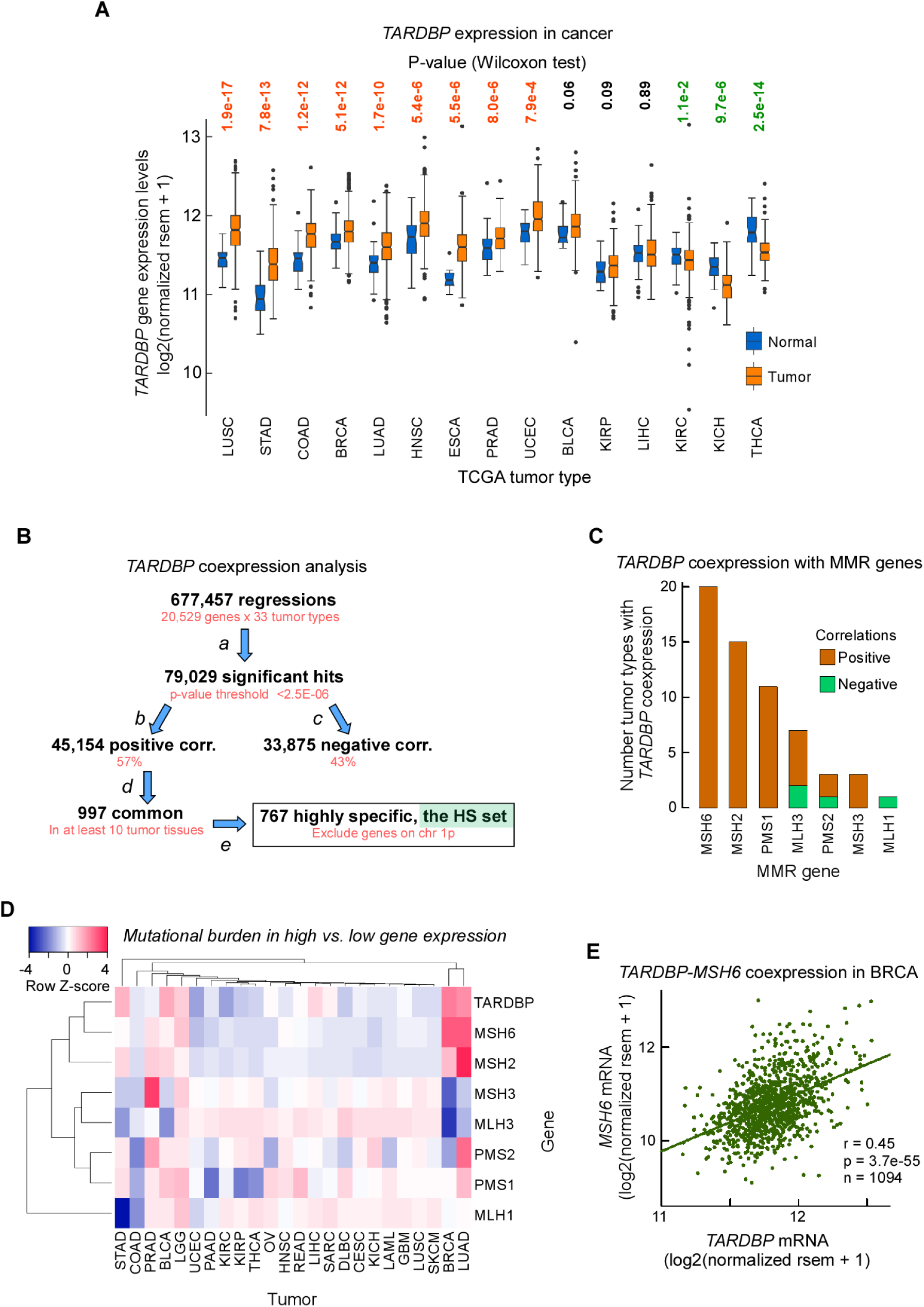
TARDB is Overexpressed in Multiple Tumor Types and Coexpresses with MSH6/2. (**A**) Box plots of *TARDBP* gene expression in 15 TCGA tumor types and matched controls. mRNA levels were normalized rsem data from RNA-seq analyses. Only tumor types with at least 10 control samples were included. Box plots display the interquartile range (IQR) from Q1 to Q3 (25–75% percentiles), median (center line), whiskers extending from Q1–1.5*IQR to Q3 + 1.5*IQR and outliers (dots). (**B**) Workflow to assess genes coexpressed with *TARDBP* in cancer. *Top*, initial screen of linear regressions of rsem normalized RNA-seq data between *TARDBP* and each of the mapped transcripts in 33 TCGA tumor types. a) filter correlations by regression coefficients with a p-value <2.5E-6, which represents an approximate Bonferroni correction for multiple comparisons (p-value/N). b and c) separate positive correlations from negative correlations. d) select genes whose correlation with *TARDBP* is observed in at least 10 different tumor types. e) filter out genes residing on the same chromosomal arm as *TARDBP* since coexpression may arise from chromosomal amplification. (**C**) Bar plot of the number of TCGA tumor types for which a significant correlation for the coexpression between *TARDBP* and MMR genes was found, from panel B (b,c). (**D**) Heat map showing mutation burden (single base substitutions) in high versus low expression of listed genes according to tumor type. (**E**) Dot plot and linear regression for the MMR gene (*MSH6*) most strongly coexpresed with *TARDBP* and tumor type (BRCA), from panel B (e).

Having found that TDP43 expression is generally dysregulated in cancer, we then assessed if the expression of TDP43 would correlate with patient survival by dividing the patients for each tumor type into two groups, a g_high group where TDP43 mRNA levels were above the population mean, and a g_low group, below the mean. A comprehensive analysis of hazard ratios in all 33 TCGA tumor types revealed that in four types of tumors, i.e. sarcoma, liver carcinoma, adrenocortical carcinoma, and low-grade glioma, patients with high TDP43 expression levels (the g_high group) would be associated with high risk of poor outcome (Supplemental **Fig. S5A**). Kaplan-Meier survival curves of this data indicated that patients with sarcoma incurred the most dramatic association between gene expression and survival, with the g_high group incurring a ∼2-fold increase in poor prognosis relative to the g_low group (Supplemental **Fig. S5B**). Therefore, assuming that *TARDB* expression correlates with TDP43 levels, our data suggest that TDP43 is an informative biomarker for some types of tumors.

Next, we developed a protocol to identify all genes coexpressed with TDP43 in cancer. From an initial screen of 677,457 linear regressions between RNA-seq data of *TARDBP* and ∼20,000 genes in each of the 33 TCGA tumor types, ∼79,000 regressions passed our threshold set at a p-value <2.5E-6 (accounting for Bonferroni correction for multiple comparisons) for significant coexpression, 57% being positive and 43% being negative associations (**Fig. 7B**). After filtering for genes positively coexpressed with *TARDBP* in at least 10 tumor types and excluding those residing on the same chromosomal arm as *TARDBP*, our algorithm returned a total of 767 highly specific genes. The strongest correlation was observed for *PNN* in acute myeloid leukemia, which displayed a regression coefficient of 0.81 and a p-value of 1.8E-41 (Supplemental **Fig. S5C**). Importantly, for the MMR genes, MSH6 was co-expressed with TDP43 in 20 tumor types, followed by MSH2 in 15 tumor types and PMS1 in 11 tumor types (**Fig. 7C**); interestingly, MSH3, whose expression correlated with that of TDP43 in only a few tumor types using our threshold, displayed the strongest association among MMR genes, with a regression coefficient of 0.78 and a p-value of 2.2E-17 in uveal melanoma (Supplemental **Fig. S5D**). From this analysis, we conclude that MSH2 and MSH6, which code for the MutSα complex for the repair of small loops and base mismatches, are strongly coexpressed with TDP43 in cancer.

Given the role of the MMR pathway in repairing DNA and the coexpression of *TARDBP* with some MMR components, we asked if there would be differences in mutational loads in cancer between the g_high and g_low groups, for both *TARDBP* and MMR genes. For *TARDBP* the strongest differences were observed in invasive breast cancer, lung adenocarcinoma, bladder urothelial carcinoma, stomach adenocarcinoma, low-grade glioma, and liver hepatocellular carcinoma, in which the g_high group incurred more mutations than the g_low group. In two additional tumors of the kidney and endometrium the g_low group displayed fewer mutations than g_high (Supplemental **Fig. S5**). For the MMR genes, hierarchical clustering placed *MSH6* in the same branch as *TARDBP* and near *MSH2*, with tumors of the breast and lung characterized by strong increases in mutational burden in patients overexpressing *TARDBP*, *MSH6* and *MSH2* (**Fig. 7D**). As anticipated, *TARDBP* was strongly coexpressed with *MSH6* and *MSH2* in invasive breast cancer and lung adenocarcinoma, with regression coefficients ranging from 0.35 to 0.47 and p-values from 7.56E-17 to 6.78E-62 (**Fig. 7E**).

Finally, to further investigate the type of genes coexpressed with TDP43 (the 767 HS set, **Fig. 7B**), we conducted a gene functional annotation analysis, which relies on the extent of overlap between a given set of genes and curated databases. The 767 HS gene set was strongly overrepresented in genes whose products undergo SUMOylation (**Fig. 8A**), a post-translational modification that plays a crucial role in nuclear transport, DNA replication and repair, mitosis, and signal transduction, and which is associated with poor outcome in cancer (76). As predicted, other types of genes overrepresented in the 767 HS set were genes involved in the cell cycle, mitosis, splicing, kinetochore formation, DNA damage and repair, and DNA replication (**Fig. 8A**). Thus, our analysis uncovered a link between TDP43 overexpression and the concomitant overexpression of genes whose processes are at the core of tumor growth. With a z-score of 2.75, the association between TDP43 and MMR was among the strongest also in normal human tissues, ranking fifth, as predicted by KEGG pathways, only preceded by splicing, RNA transport, DNA replication and cell cycle. As per individual genes, at a z-score >5 for predicted biological processes, both for *MSH2*, *MSH6* and *MLH1*, and most of the top genes coexpressed with TDP43, the strongest association was with DNA unwinding involved in DNA replication (**Fig. 8B**). In summary, our in-silico analysis confirms a strong association of TDP43 with the MMR pathway, which is seen to occur in the context of cell cycle regulation and protection of genome fidelity in a normal cell, but which represent processes that are strongly activated in cancer to achieve uncontrolled cell division and high genome instability.

**Figure 8.**
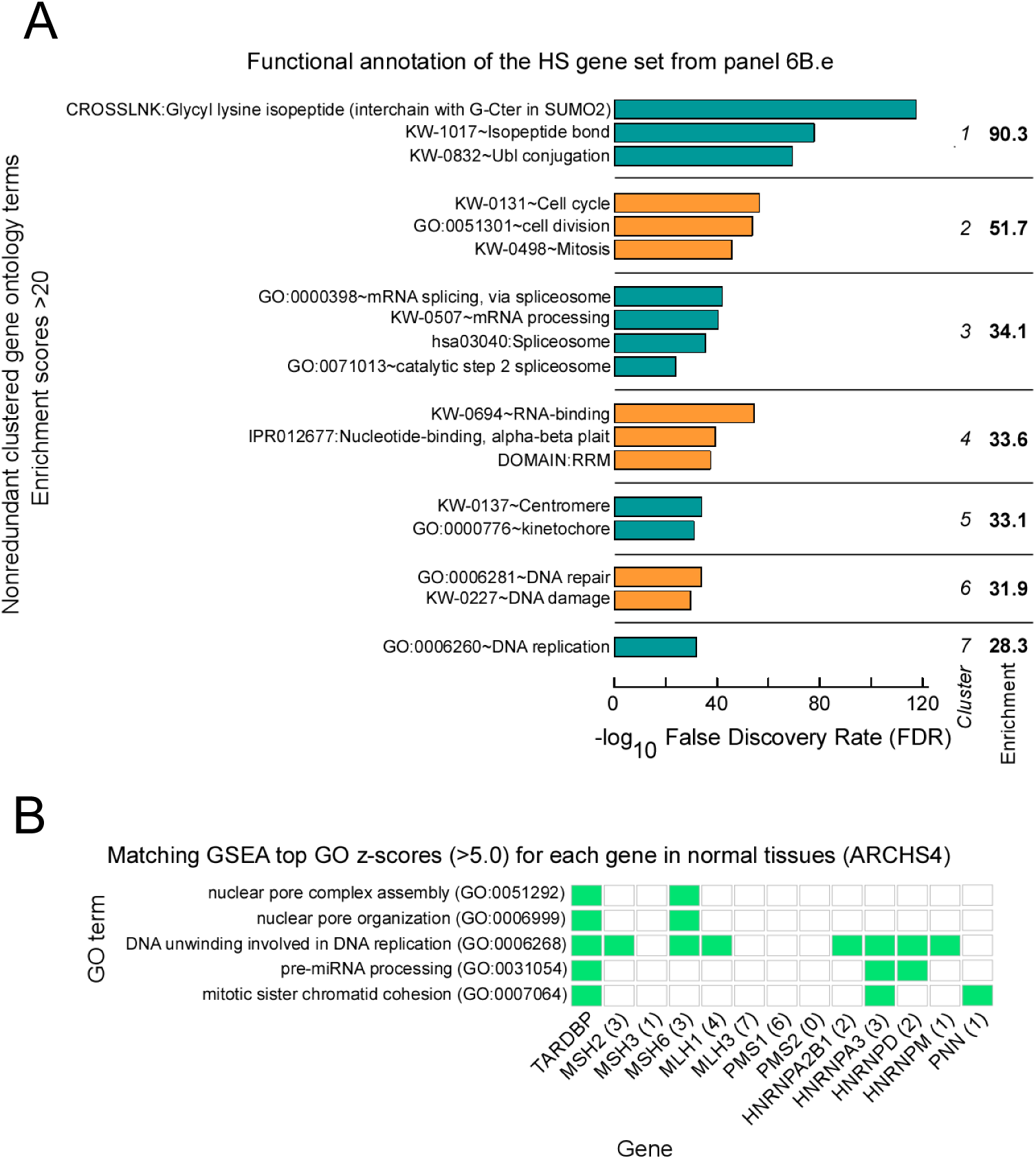
Genes Co-expressed with *TARDBP* are at the Core of Cell Cycle and DNA Repair. (**A**) Gene Set Enrichment Analysis (GSEA) of genes highly coexpressed with *TARDBP*. Only enrichment scores >20 and the ontology terms with the strongest enrichment score for redundant ontology terms were displayed. (**B**) Check box for predicted biological processes (GO) with z-scores >5 for *TARDBP*, and *TARDBP*-matching GO terms with z-scores >5 for MMR genes and HNRNP genes highly coexpressed with *TARDBP* at levels above that of the *PNN* gene (Pearson’s coefficient >0.63 in human normal tissues). In parenthesis the number of GO terms with z-scores >5.

## Discussion

Two major age-related disease areas with critical grand challenges for biology and medicine are etiology and intervention for neurodegeneration and cancer. MMR balance has been implicated in both processes. This observation raises the question as to how MMR balance is dynamically regulated in cells. As TDP43 has been linked to DDR gene expression and found to have a direct role in DSB repair (19), we reasoned that TDP43 may be a key factor for actively regulating MMR balance. By employing the RT2-Profiler assay to analyze HEK293 RNA extracts, we discovered a consistent decrease in the expression of several MMR genes following TDP43 KD. This phenomenon was not limited to a specific cell type; both NPCs and iMNs also exhibited suppressed MMR function upon TDP43 KD. Having established this clear phenotype, we sought to uncover the potential mechanisms.

TDP43 is widely recognized for its involvement in RNA processing. Likewise, its pathological mislocalization has been linked to detrimental transcriptomic alterations in both experimental and post-mortem ALS tissues (1, 8, 59, 77–79). Investigating TDP43’s effect on MMR transcripts, we utilized *in silico* tools and manual identification of UG-rich repeat regions to predict TDP43 binding sites on *MLH1* and *MSH6* transcripts (80). We chose these genes for their clear binding potential and consistent co-expression with TDP43. We then employed end-point RT-PCR with primers designed to amplify the identified regions to ascertain whether the loss of TDP43 led to the inclusion of intron sequences. Based on the sizes of amplicon products, we conclude that TDP43 KD negatively affects the transcriptional processing of these genes. Furthermore, we observed this impact also occurred in cell models expressing different ALS-associated TDP43 mutations, as well as in RNA samples isolated from Guamanian ALS tissues. Significantly, these results align with previous reports identifying TDP43-dependent splicing of ALS disease-related genes (1, 8, 59, 61, 77, 79). Delving deeper into the consequences of mis-splicing and considering the potential of decreased transcript stability resulting from it, we tested the effect of TDP43 on the stability of MLH1 and MSH6 transcripts (81, 82). Utilizing α-amanitin as an RNA polymerase II inhibitor, we conducted an RNA stability experiment that measured transcript abundance over time. Using this approach, we confirmed that TDP43 KD caused an accelerated decrease in MLH1 and MSH6 mRNA levels, suggesting a mechanism where TDP43 modulates MMR expression through modulating transcript stability. Conversely, we observed the opposite effect on levels of TDP43 itself following its autoregulatory function (1).

Notably, our results highlighted the bidirectional nature of TDP43’s influence on MMR expression. Differentiated cells showed a decrease in MMR protein levels, which was reversible by WT TDP43 OE, which not only declined MMR expression in nondividing cells but also increased MMR expression to levels higher than those observed in dividing controls. Significantly, we also observed that MMR expression decreased with each stage of cellular differentiation of iPSCs into NSCs and then into motor neurons. To determine whether this effect was relevant to neurodegenerative disease, we measured MMR expression in the brain tissue of two separate ALS mouse models. The first utilized low overexpression of murine Tdp43, while the second utilized a Tdp43ΔNLS knock-in mutation without a change in copy number (70). Both models exhibited increased MMR expression, albeit with slight differences. The fact that our Tdp43ΔNLS mouse model exhibited this phenotype was intriguing. We suspect the mislocalization of TDP43 in these cells induces a relative increase in TDP43 via its autoregulatory feedback mechanism (1). This, in turn, increases MMR expression similar to that observed in the TDP43 overexpression model. We also repeated these tests in a third, more aggressive mouse model of ALS that expresses very high levels of human TDP43 and severe early onset of disease. Unexpectedly, we did not observe a widespread increase in MMR expression. We reason that this may be an artifact of the sudden and severe cellular pathology exhibited by this model, which causes rapid apoptosis of affected cells. Together, these models complemented each other by capturing distinct but overlapping pathological features of TDP-43 dysfunction, isolated mislocalization versus overexpression-induced mislocalization, thus strengthening the translational relevance and robustness of our findings. Notably, we observed sex-specific differences in MSH2 levels in the Prp mouse model. While a detailed investigation of the underlying mechanisms is beyond the scope of the current study, previous work has shown sex differences in cancer risk associated with MSH2 mutations, such as a slightly earlier onset in males; the overall expression or methylation status of MSH2 has not been consistently shown to be sex dependent (83, 84). Our data, however, indicate female-specific MSH2 OE in the context of TDP-43 pathology, which could suggest novel sex-linked regulatory mechanisms in the brain’s response to neurodegenerative stresses, warranting further investigations on this crucial aspect of genetic mutation regulation. Finally, we measured the expression of MMR protein from CNS tissues of Guamanian ALS-affected patients. Despite limited technical hindrances, we reliably show that at least one MMR protein, MLH1, is significantly higher in the insoluble protein fractions from ALS tissues than in controls. Taken together, these observations highlight the strong influence of TDP43 on MMR expression as it reverses natural MMR downregulation during cellular differentiation while increasing expression in models and tissues of ALS-TDP43.

When considering how TDP43 may affect these changes in MMR expression, two potential mechanisms may be considered. First, our findings show that TDP43 modulates transcriptional splicing and stability of MLH1 and MSH6 transcripts, suggesting that other MMR factors may be regulated similarly. While we did not explicitly study how TDP43 influences transcript splicing and stability of every MMR gene, TDP43 is known to interact with hundreds of different transcripts (59). The true scope of its influence is likely much larger. Therefore, it is plausible that TDP43 may also directly affect other MMR transcripts in a manner not yet described. For this study, the demonstration of how TDP43 affects the splicing and stability of MLH1 and MSH6 highlights representative mechanisms. Furthermore, given the interdependent functions of the MMR proteins, it is reasonable to conclude that the dysregulation of one protein is likely to affect the overall functionality of the MMR system; for example, absence or mutation of a single MMR protein, MSH2, is sufficient to cause colorectal cancer. Second, TDP43 overexpression likely causes network-level adjustments to the cellular transcriptome, leading to a multitude of changes, of which increased MMR expression is a part. Our bioinformatics experiments showing the functional annotation of genes enriched by TDP43 overexpression support this notion. Several of the identified proteins were closely linked to cell cycle control, mitosis, splicing, kinetochore formation, DNA damage/repair, and DNA replication. This aligns with other reports on TDP43’s contribution toward maintaining cancer cell stemness and cellular differentiation (85, 86). There are also implications for neurodegeneration. Specifically, Alzheimer’s disease-affected neurons adopt an altered transcriptomic profile characterized by markers of cell cycle alteration and dedifferentiation from the neuronal state (87). In both cases, TDP43 pathology is associated with disease-relevant alterations in the cellular transcriptome. Exhaustive dissection of the myriad consequences of these changes is still lacking. However, our data indicates MMR dysregulation may be a part of this complex system of events.

To better understand how MMR dysregulation may contribute to TDP43-linked disease processes, we utilized a previously reported neuronal cell model of mislocalized TDP43 expression (88). In this model, as in our Tdp43ΔNLS mouse model, the expression of mislocalized TDP43 increased MMR expression. To test the contribution of MMR, we chose to isolate its effect via KD of MSH2; this effectively diminishes MMR damage recognition capability and negatively affects the expression and activity of other MMR factors (25, 89, 90). Using the sensitive technique of alkaline single-cell gel electrophoresis, we observed that MSH2 KD partially rescued Tdp43ΔNLS-induced DNA damage. Furthermore, we also showed MSH2 KD improved cellular capacity to repair oxidative DNA damage by glucose oxidase with concomitant decreases in markers of DDR signaling activation following a 6 h recovery period.

These findings may have important implications for both cancer and neurodegenerative pathologies. From a neuronal perspective, our results showing MMR enhancing Tdp43ΔNLS-induced DNA damage, DDR signaling, and apoptosis are novel. They also align with existing reports of increased DDR signaling and DNA damage observed in neurodegenerative disease (91–93). Exactly how MMR exerts this effect is poorly understood. Multiple reports suggest that MMR activity in nondividing cells is likely mutagenic, possibly due to a lack of a strand discrimination signal (48, 49). This alone could be a direct source of increased DNA damage and DDR signaling. The response of MMR proteins to unique nucleic acid structures associated with neurodegenerative diseases, such as R-Loops and trinucleotide repeats (TNR) mutations, is also poorly understood. Exciting new work has linked MMR to the progression of TNR diseases such as Huntington’s disease (94), myotonic dystrophy (MD), and fragile X syndrome (FXS), among others (51, 53–56, 94, 95). MLH1, for example, was shown to play a direct role in the expansion of certain TNR mutations, leading to disease progression in a mouse model of HD (52). R-loops are another unique structure associated with ALS and other neurodegenerative diseases. These hybrid structures of RNA and DNA exist transiently at sites of heavy transcription and have been associated with TDP43 dysfunction, TNR expansion, and C9ORF72 ALS progression (96–101). Interestingly, MMR activity has also been linked to sites of heavy transcription (23, 74, 102), highlighting the importance of understanding how MMR may interact with R-loop structures.

From a cancer perspective, the loss of MMR is a well-known step along the path of carcinogenesis, but its loss is equally important for chemotherapeutic cancer cell killing. This second point is based on MMR-mediated DDR and apoptotic signaling (31, 32, 40, 41). It has been shown in multiple cancers of the endometrium, colon, and brain that dMMR impairs the cell-killing ability of chemotherapy, especially for platinum-based and alkylating agents (36–39). Surprisingly, overexpressing MMR in dividing cells has also been linked to a mutator phenotype and DDR activation (42–44, 48, 103). For example, *in vitro* overexpression of either MLH1 or PMS2 disrupted normal MMR activity, leading to increased mutations (42, 43). Our computational analyses revealed a strong association of TDP43 with the MMR pathway, with clear trends of TDP43 and MMR co-expression. We furthermore found that TDP43 is often upregulated in cancer and confers a poor prognosis. These observations align with previous reports associating TDP43 with enhanced tumor progression and treatment resistance (104, 105). Taken together, our results reveal new mechanistic links for TDP43 with MMR and cancer pathology.

Our study is not without limitations. There remains a need to fully characterize how TDP43 affects the transcriptional splicing and stability of all MMR factors. Additionally, our exploration of MMR’s contribution to DNA damage and TDP43 pathology was but the first step toward understanding the full consequences of this molecular relationship. Finally, our exploration of the TDP43-MMR relationship in cancer would benefit from in vitro testing of hypotheses generated from *in silico* analyses.

Currently, our collective data uncover a fundamental finding of TDP43-mediated control of MMR expression that extends across multiple cell models, animal models, and disease conditions. Our data, in combination with previous reports, present TDP43-mediated regulation of MMR as part of a shared cellular program with critical implications in both neurodegeneration and cancer. We anticipate that this discovery will serve as a foundation for future work testing how this relationship may potentially be leveraged to treat disease processes ranging from TNR mutations in HD to improved targeting of cancer stem cells. As the medical establishment must increasingly confront the harsh realities of caring for an aging population, the importance of understanding the underlying molecular mechanisms shared by these two major age-related disease classes with the potential for positive interventions should not be understated.

## Acknowledgements

This research is primarily supported by the National Institute of Neurological Disorders and Stroke (NINDS) and the National Institute on Aging of the National Institutes of Health (NIH) under award number RF1NS112719 to M.L.H. Research in the Hegde laboratory is also supported by the NIH awards R01NS088645, R03AG064266, and R01NS094535, as well as the Sherman Foundation Parkinson’s Disease Research Challenge Fund and the Houston Methodist Research Institute’s internal funds. M.L.H also acknowledges Everett E. and Randee K. Bernal for their support via the Centennial Endowed Chair of Neurological Institute. Efforts by J.A.T and A.B are supported by NIH P01 CA092584 and R35 CA220430, Cancer Prevention and Research Institute of Texas RP180813 and Robert A. Welch Chemistry Chair (G-0010). This work used HPC resources at the Texas Advanced Computing Center (TACC) at The University of Texas at Austin (URL: http://www.tacc.utexas.edu). The authors thank Dr. Anna Dodson at Houston Methodist Research Institute (Houston, TX) for assistance with document editing and other members of the Hegde laboratory at Houston Methodist.

## Author Contributions

V.E.P. performed experiments with assistance from S.R., J.M., M.K., V.H.M.R, and V.V. V.E.P. and J.M. conducted data analysis with interpretation and wrote the manuscript. S.M., R.G., Z.X., and J.A.T. provided valuable tissues, intellectual insights, analysis, and manuscript feedback. A.B. performed bioinformatics analysis and wrote results for **Figs. 6** and **7**. M.L.H. supervised the study and prepared the final manuscript. All authors participated in the discussion and provided valuable feedback on the manuscript.

## Data Availability Statement

Raw data for experiments in this study can be provided upon reasonable request to the corresponding author. A copy of normalized TCGA RNA-seq data is available at https://doi.org/10.5281/zenodo.7885656. Source codes used for the TCGA bioinformatic analyses are available at https://doi.org/10.5281/zenodo.7874703. C++ codes for t-test and linear regression are available at https://github.com/abacolla.

