## Supplementary material for "RNA/DNA Binding Protein TDP43 Regulates DNA Mismatch Repair Genes with Implications for Genome Stability"

Muralidhar L Hegde, PhD
Professor, Department of Neurosurgery
Everett E. and Randee K. Bernal Centennial Endowed Chair of Neurological Institute
Director, Division of DNA Repair Research, Center for Neuroregeneration
Houston Methodist Research Institute
6670 Bertner Ave, RI-11-112

Supplementary Material associated with this article includes six Tables and five Figures.

**Supplemental Table S1:** List of genes affected by TDP-43 downregulation in HEK-293 cells on $RT^2$ profiler assay.

| | Gene name | Gene description | Fold change Si.TDP43 vs si.Control | DNA repair pathway |
|---|---|---|---|---|
| 1 | MLH1 | MutL homolog 1, colon cancer, nonpolyposis type 2 (E. coli) | 0.1319 | Mis-match Repair |
| 2 | MLH3 | MutL homolog 3 (E. coli) | 0.9783 | |
| 3 | MSH2 | MutS homolog 2, colon cancer, nonpolyposis type 1 (E. coli) | 0.5017 | |
| 4 | MSH3 | MutS homolog 3 (E. coli) | 0.215 | |
| 5 | MSH4 | MutS homolog 4 (E. coli) | 2.2542 | |
| 6 | MSH5 | MutS homolog 5 (E. coli) | 1.0374 | |
| 7 | MSH6 | MutS homolog 6 (E. coli) | 0.2628 | |
| 8 | PMS1 | PMS1 postmeiotic segregation increased 1 (S. cerevisiae) | 0.411 | |
| 9 | PMS2 | PMS2 postmeiotic segregation increased 2 (S. cerevisiae) | 0.1976 | |
| 10 | POLD3 | Polymerase (DNA-directed), delta 3, accessory subunit | 0.0249 | |
| 11 | TREX1 | Three prime repair exonuclease 1 | 1.8804 | |
| 1 | ATXN3 | Ataxin 3 | 4.2193 | Other pathways |
| 2 | BRIP1 | BRCA1 interacting protein C-terminal helicase 1 | 1.161 | |
| 3 | CCNH | Cyclin H | 0.6207 | |
| 4 | CDK7 | Cyclin-dependent kinase 7 | 0.5785 | |
| 5 | DDB1 | Damage-specific DNA binding protein 1, 127kDa | 0.3256 | |
| 6 | DDB2 | Damage-specific DNA binding protein 2, 48kDa | 0.64 | |
| 7 | ERCC1 | Excision repair cross-complementing rodent repair deficiency, complementation group 1 (includes overlapping antisense sequence) | 0.3252 | |
| 8 | ERCC2 | Excision repair cross-complementing rodent repair deficiency, complementation group 2 | 0.1092 | |
| 9 | ERCC3 | Excision repair cross-complementing rodent repair deficiency, complementation group 3 (xeroderma pigmentosum group B complementing) | 0.67 | |
| 10 | ERCC4 | Excision repair cross-complementing rodent repair deficiency, complementation group 4 | 0.6857 | |
| 11 | ERCC5 | Excision repair cross-complementing rodent repair deficiency, complementation group 5 | 2.2542 | |
| 12 | ERCC6 | Excision repair cross-complementing rodent repair deficiency, complementation group 6 | 2.2542 | |
| 13 | ERCC8 | Excision repair cross-complementing rodent repair deficiency, complementation group 8 | 0.2492 | |
| 14 | LIG1 | Ligase I, DNA, ATP-dependent | 0.4863 | |
| 15 | MMS19 | MMS19 nucleotide excision repair homolog (S. cerevisiae) | 1.178 | |
| 16 | PNKP | Polynucleotide kinase 3'-phosphatase | 2.2542 | |
| 17 | POLL | Polymerase (DNA directed), lambda | 0.406 | |
| 18 | RAD23A | RAD23 homolog A (S. cerevisiae) | 2.3739 | |
| 19 | RAD23B | RAD23 homolog B (S. cerevisiae) | 0.2463 | |
| 20 | RPA1 | Replication protein A1, 70kDa | 0.3135 | |
| 21 | RPA3 | Replication protein A3, 14kDa | 2.0581 | |
| 22 | SLK | STE20-like kinase | 0.8157 | |
| 23 | XAB2 | XPA binding protein 2 | 1.3035 | |
| 24 | XPA | Xeroderma pigmentosum, complementation group A | 1.0014 | |
| 25 | XPC | Xeroderma pigmentosum, complementation group C | 1.2946 | |

| | | | | |
|---|---|---|---|---|
| 1 | APEX1 | APEX nuclease (multifunctional DNA repair enzyme) 1 | 0.3295 | Base Excision Repair |
| 2 | APEX2 | APEX nuclease (apurinic/apyrimidinic endonuclease) 2 | 0.0327 | |
| 3 | CCNO | Cyclin O | 2.2542 | |
| 4 | LIG3 | Ligase III, DNA, ATP-dependent | 0.7082 | |
| 5 | MPG | N-methylpurine-DNA glycosylase | 2.2542 | |
| 6 | MUTYH | MutY homolog (E. coli) | 0.9966 | |
| 7 | NEIL1 | Nei endonuclease VIII-like 1 (E. coli) | 16.6977 | |
| 8 | NEIL2 | Nei endonuclease VIII-like 2 (E. coli) | 2.2542 | |
| 9 | NEIL3 | Nei endonuclease VIII-like 3 (E. coli) | 1.2653 | |
| 10 | NTHL1 | Nth endonuclease III-like 1 (E. coli) | 2.5263 | |
| 11 | OGG1 | 8-oxoguanine DNA glycosylase | 0.6781 | |
| 12 | PARP1 | Poly (ADP-ribose) polymerase 1 | 0.3292 | |
| 13 | PARP2 | Poly (ADP-ribose) polymerase 2 | 0.6526 | |
| 14 | PARP3 | Poly (ADP-ribose) polymerase family, member 3 | 4.3363 | |
| 15 | POLB | Polymerase (DNA directed), beta | 3.2099 | |
| 16 | SMUG1 | Single-strand-selective monofunctional uracil-DNA glycosylase 1 | 0.0486 | |
| 17 | TDG | Thymine-DNA glycosylase | 0.14 | |
| 18 | UNG | Uracil-DNA glycosylase | 0.4187 | |
| 19 | XRCC1 | X-ray repair complementing defective repair in Chinese hamster cells 1 | 0.8819 | |
| 1 | BRCA1 | Breast cancer 1, early onset | 0.6827 | Double-strand break |
| 2 | BRCA2 | Breast cancer 2, early onset | 0.2023 | |
| 3 | DMC1 | DMC1 dosage suppressor of mck1 homolog, meiosis-specific homologous recombination (yeast) | 2.2542 | |
| 4 | FEN1 | Flap structure-specific endonuclease 1 | 0.527 | |
| 5 | LIG4 | Ligase IV, DNA, ATP-dependent | 0.8476 | |
| 6 | MRE11A | MRE11 meiotic recombination 11 homolog A (S. cerevisiae) | 0.5124 | |
| 7 | PRKDC | Protein kinase, DNA-activated, catalytic polypeptide | 0.3737 | |
| 8 | RAD21 | RAD21 homolog (S. pombe) | 0.8458 | |
| 9 | RAD50 | RAD50 homolog (S. cerevisiae) | 0.3231 | |
| 10 | RAD51 | RAD51 homolog (S. cerevisiae) | 0.3289 | |
| 11 | RAD51B | RAD51 homolog B (S. cerevisiae) | 1.3087 | |
| 12 | RAD51C | RAD51 homolog C (S. cerevisiae) | 0.4217 | |
| 13 | RAD51D | RAD51 homolog D (S. cerevisiae) | 1.1946 | |
| 14 | RAD52 | RAD52 homolog (S. cerevisiae) | 0.2345 | |
| 15 | RAD54L | RAD54-like (S. cerevisiae) | 0.3607 | |
| 16 | XRCC2 | X-ray repair complementing defective repair in Chinese hamster cells 2 | 1.0313 | |
| 17 | XRCC3 | X-ray repair complementing defective repair in Chinese hamster cells 3 | 2.2542 | |
| 18 | XRCC4 | X-ray repair complementing defective repair in Chinese hamster cells 4 | 0.3446 | |
| 19 | XRCC5 | X-ray repair complementing defective repair in Chinese hamster cells 5 (double-strand-break rejoining) | 1.5425 | |
| 20 | XRCC6 | X-ray repair complementing defective repair in Chinese hamster cells 6 | 0.8264 | |
| 1 | ATM | Ataxia telangiectasia mutated | 2.2542 | Other |
| 2 | ATR | Ataxia telangiectasia and Rad3 related | 2.395 | |
| 3 | EXO1 | Exonuclease 1 | 0.9994 | |
| 4 | MGMT | O-6-methylguanine-DNA methyltransferase | 1.3726 | |
| 5 | RFC1 | Replication factor C (activator 1) 1, 145kDa | 0.3373 | |
| 6 | TOP3A | Topoisomerase (DNA) III alpha | 1.3096 | |
| 7 | TOP3B | Topoisomerase (DNA) III beta | 2.1114 | |
| 8 | XRCC6BP1 | XRCC6 binding protein 1 | 1.4236 | |

**Supplemental Table S2:** List of primer pairs of MMR-associated and housekeeping genes for qRT-PCR assay.

| Primer | Forward | Reverse |
|---|---|---|
| MLH1 | TTCACCCAGACTTTGCTACCAGG | GAAGTAGGTCCTCTCTTCTCTGACAT |
| MSH2 | GGTATTTGGCATATAAGGCTTCTCCTG | CATGTCTCCAGCAGTCTCTCCTC |
| MSH3 | TGGCAACTCTGAGCCAAAGAAATG | GCATTCTTTGCGTGTAGAAGACTGATAT |
| MSH6 | TCGCAGTGTTGGATGTTTTACTGTG | TGTCTCATAAGCGTAGACTTGCCC |
| PMS2 | TGCACTGAGCGATGTCACCATTTC | GAAAATAACTGCTGCACGCTGACT |
| TDP43 | GTGTGGGCTTCGCTACAGG | CAACATACACCAGATTTCCCCAG |
| HPRT | CCTGGCGTCGTGATTAGTGAT | AGACGTTCAGTCCTGTCCATAA |
| 18S rRNA | GACTGTCTCGCCGGTGTC | GGAGAGCCGGAACGTCGA |

**Supplemental Table S3:** List of primer pairs for intron-exon inclusion/exclusion assay of MMR-associated genes by RT-PCR.

| Primer | Forward | Reverse |
|---|---|---|
| MLH1 Ex17-Ex18 | CTATGTGCCCCCTTTGGAGG | CCGGATGGAATAGAACATAGCG |
| MSH3 Ex14-Ex15 | ACCCAAGAGTTCTTCTTGATTGTCA | ACTGTCACATATTGTGCAGAAGGA |
| MSH6 Ex11-Ex12 | CCCAGCCAGGAGACTATTACG | GCAATTTATGGACAGCTTCAGCA |
| MSH6 Ex8-Ex9 | GAGTGTTTACTAGACTTGGTGCCTCAG | TGCTGTTGCATGCATGAGTATGC |
| PMS2 Ex2-Ex3 | CCTGCTAAGGCCATCAAACCTATT | CTTCGAAGTTTTCTTCTTCTACCCCAC |
| GAPDH | GGAGCGAGATCCCTCCAAAAT | GGCTGTTGTCATACTTCTCATGG |

**Supplemental Table S4:** List of primers for mini gene assay

| Primer | Forward | Reverse |
|---|---|---|
| MLH1 Ex17-Ex18 Cloning (RHC-Glo) | **GGCCGGATCC**GAAGGGAACCTGATTGGAATTA | **GGCCCTCGAG**CTGCTGGCCTGAGAGGGT |
| RHCglo primers for verification | GCTTTCCTGATGCCTTCTGG | GAGTGCTTTCCTAGCCTTCC |

**Supplemental Table S5:** Summary of murine Tdp43 interactions with MMR gene transcripts in the mouse brain.

| Gene | NCBI Ref ID | Chr# | No. of Tdp-43 binding motifs | Exon/Intron |
|---|---|---|---|---|
| Mlh1 | NC_000075.7 | 9 | 5 | Intron |
| Msh2 | NC_000083.7 | 17 | 3 | Intron |
| Msh3 | NC_000079.7 | 13 | 15 | Intron |
| Msh6 | NC_000083.7 | 17 | 0 | N/A |
| Pms2 | NC_000071.7 | 5 | 2 | Intron |

**Supplemental Table S6:** Demographics of Guamanian-ALS and non-ALS control patients' brain samples.

| Identification No. | Tissue Type | Diagnosis | Age (yr) | Gender | Ethnicity |
|---|---|---|---|---|---|
| 311 | Hippocampus, Cortex | Control | 57 | F | Chamorro |
| 3842 | Cortex | Control | 48 | M | Chamorro |
| 3855 | Hippocampus, Cortex | Control | 43 | F | Chamorro |
| 3856 | Cortex | Control | 63 | M | Chamorro |
| 314 | Hippocampus | ALS | 54 | F | Chamorro |
| 362 | Hippocampus | ALS | 59 | M | Chamorro |
| 3882 | Hippocampus, Cortex | ALS | 42 | M | Chamorro |
| 3912 | Hippocampus, Cortex | ALS | 47 | F | Chamorro |

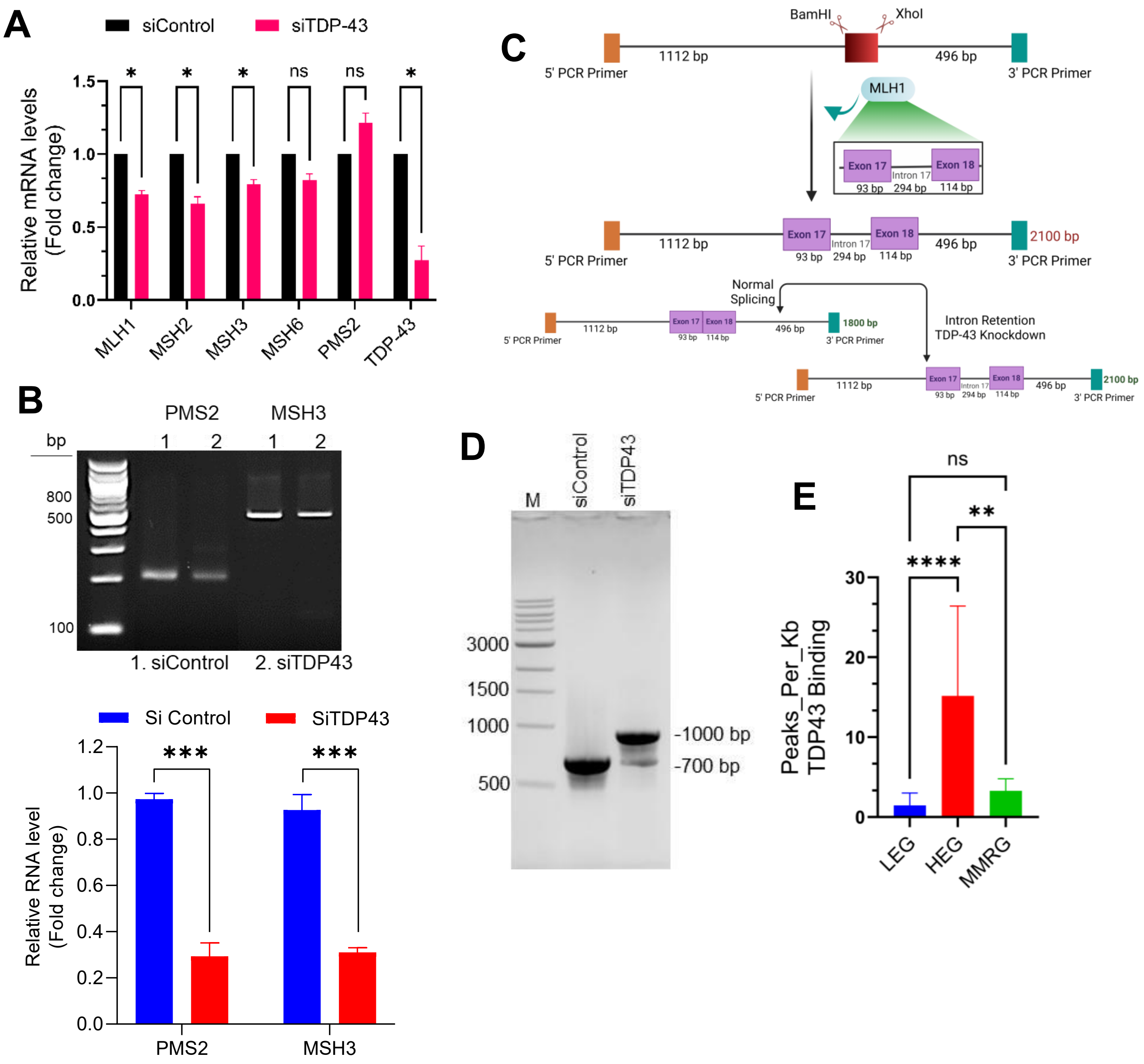


**Supplementary Figure S1 (Related to Fig. 2 and 3):**

**(A)** QRT-PCR analysis of MMR gene expression in NPSCs with or without TDP43 downregulation. (**B**) Additional tests of transcript splicing are shown using siControl or siTDP43-treated HEK-293 cells. PCR amplification and quantitation of selected regions of demonstrate loss of TDP43 did not affect splicing of PMS2 Exon 2 — Exon 3, and MSH3 Exon 14 — Exon 15. (**C-D**) A schematic of the minigene assay construct for *MLH1* gene (**C**). (**D**) A 2% agarose gel image displaying differences in PCR product sizes for *MLH1* transcripts from siControl or siTDP43-treated cells. (**E**) Bar graphs indicate a statistically significant number of murine Tdp43-bound peaks per Kb region on MMR genes (MMRG) in the mouse brain, when compared to 20 low expression (LEG) and 20 high expression genes (HEG). The groups were compared using one-way or two-way ANOVA as appropriate. The p-values are as follows: p >0.5 (ns), p <0.5 (*). Error bars indicate mean ± SEM..

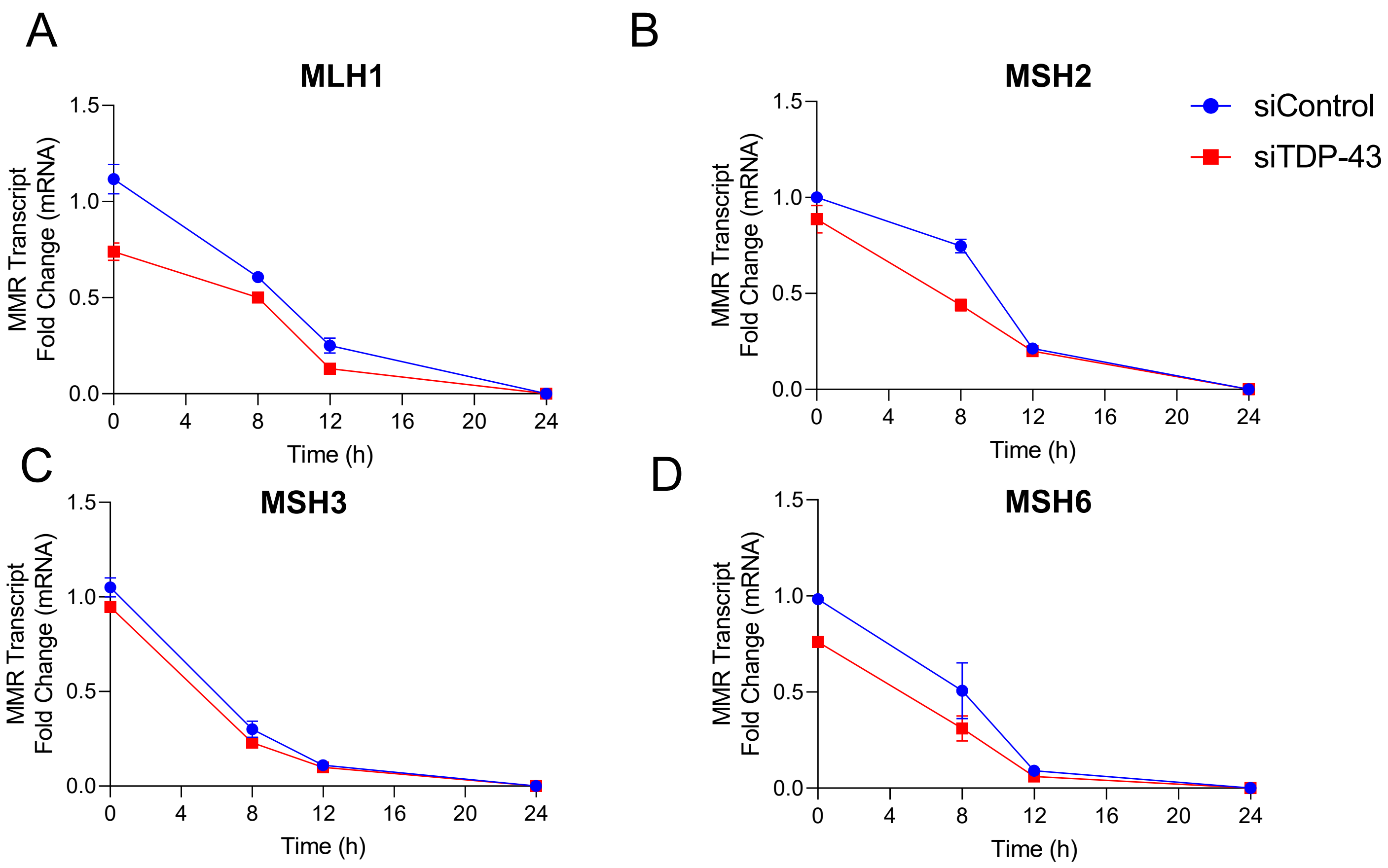


**Supplementary Figure S2 (Related to Fig. 4):**
TDP43 regulates the stability of select MLH1 and MSH6 transcripts. (**A-D**) Expressions of MMR-associated gene transcripts in relative fold changes as assessed by qRT-PCR assays for MLH1, MSH2, MSH3, and MSH6.

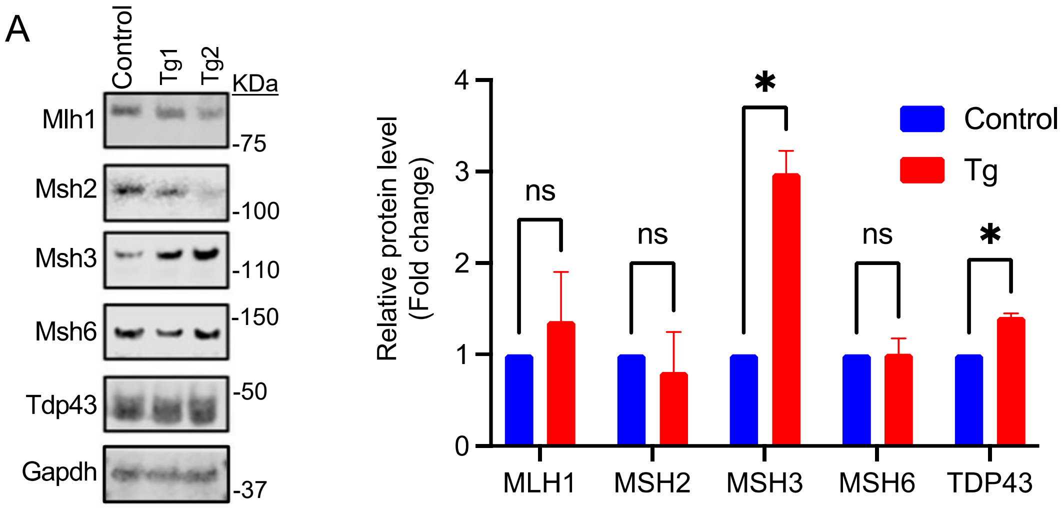


**Supplementary Figure S3 (Related to Fig. 5):**

(**A**) Immunoblot (IB) images of cortical brain lysates from Thy1 promoter-driven human TDP43 OE transgenic mouse brain samples. Histograms showing fold change in immunoblot band intensity of averaged Tg samples (red) relative to control (blue) indicate significant increases in Msh3 with trends of increasing Mlh1 expression. Significance values (p-values) are as follows: $p > 0.5$ (ns), $p < 0.5$ (*). Error bars indicate mean ± SEM from three technical replicates.

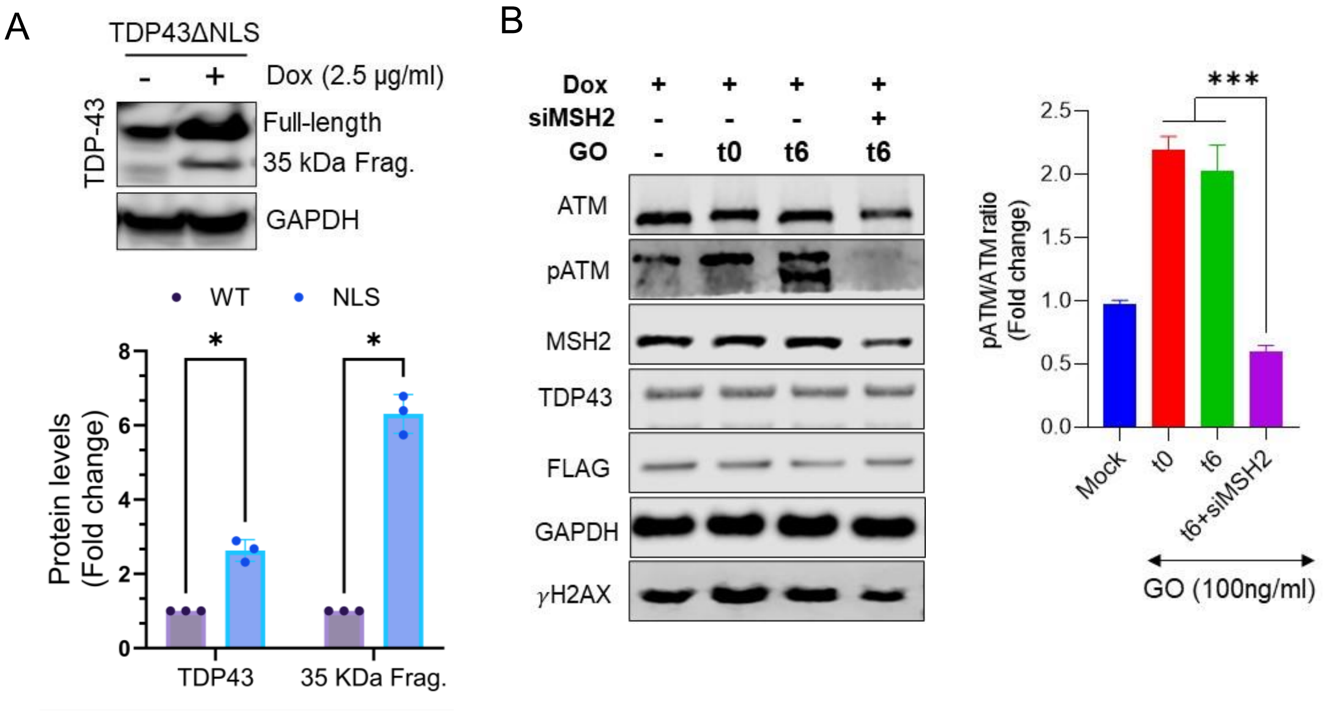


**Supplementary Figure S4 (Related to Fig. 6).**
(**A**) IB analysis of TDP43 mislocalization into the cytosol and formation of toxic 35 kDa fragment in Dox-induced or uninduced TDP43ΔNLS SH-SY5Y cell line. GAPDH served as the loading control for the cytosolic extracts. Quantification of IB band intensities for full-length and 35 kDa fragment of TDP43. (**B**) IB analysis with quantitation histograms of control and dox-inducible TDP43ΔNLS expressing cells immediately after (t0) and 6 h recovery period (t6) after 45 min exposure to glucose oxidase (GO). Quantification of GO treatment shows a significant increase in the expression of phosphorylated ATM (pATM), and H2AX ($\gamma$H2AX), which were partially rescued by MSH2 knockdown. $p < 0.001$ (***). Error bars indicate mean ± SEM from three technical replicates. The groups were compared using one-way or two-way ANOVA.

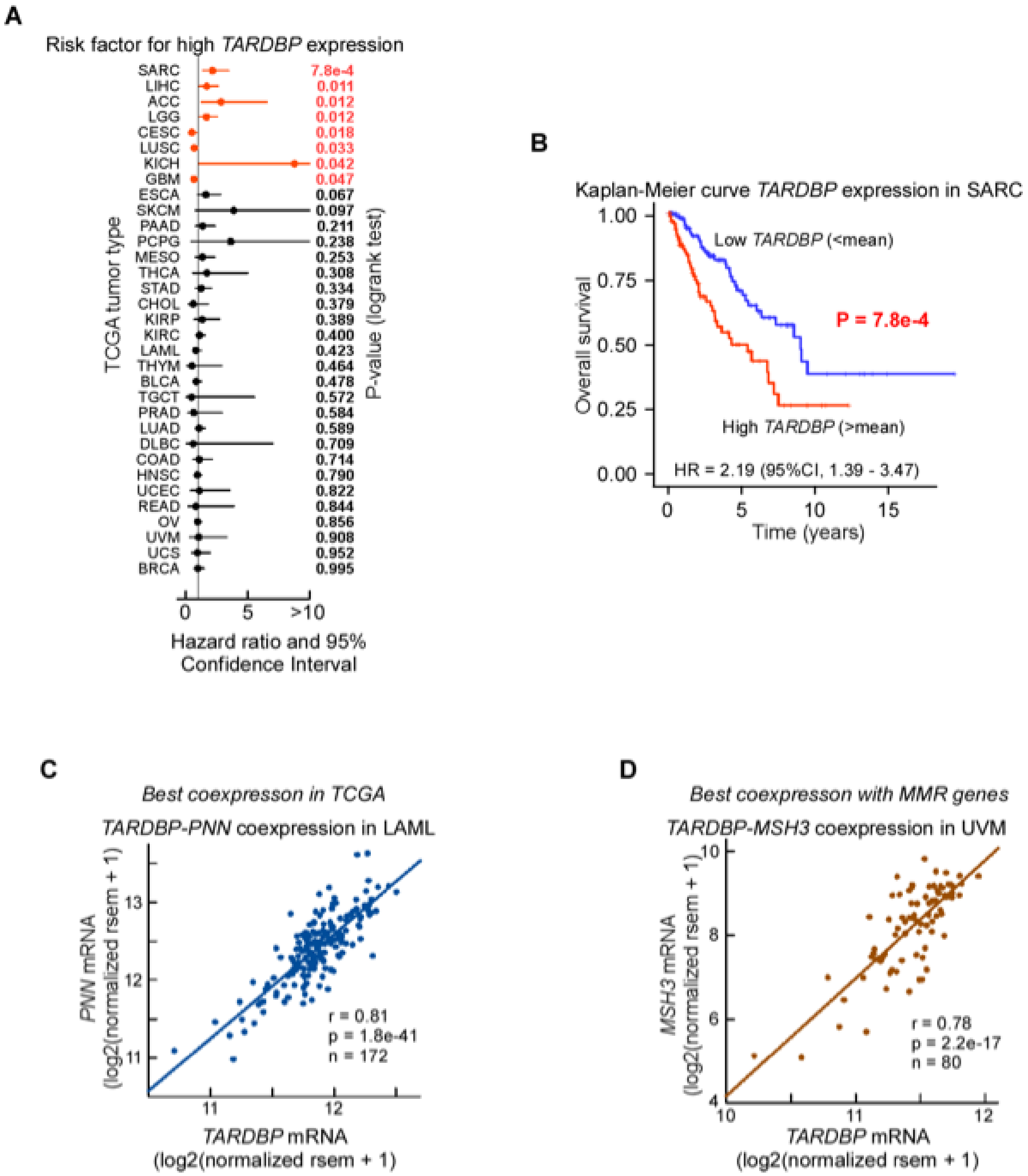


**Supplementary Figure S5 (Related to Figs. 7 and 8):**

(**A**) Hazard ratios and 95% confidence intervals (CI) sorted by P-values for patients expressing high (above population mean) versus low (below population mean) mRNA levels of *TARDBP* in all 33 TCGA tumor types. Upper CI values were truncated at 10 and were 19.32 for SKCM, 31.38 for PCPG and 71.7 for KICH. P-values from logrank tests. (**B**) Kaplan-Meier overall survival curve for patients with sarcoma from TCGA project expressing high (above population mean) versus low (below population mean) mRNA levels of *TARDBP.* P-value from logrank test. (**C**) Dot plot and linear regression for the gene (*PNN*) most strongly coexpresed with *TARDBP* and tumor type (LAML). P-value from the t-distribution. (**D**) Dot plot and linear regression for the MMR gene (*MSH3*) most strongly coexpresed with *TARDBP* and tumor type (UVM).
